# Combined effects of *Ret* coding and enhancer loss-of-function alleles cause progressive loss of inhibitory motor neurons in the enteric nervous system

**DOI:** 10.1101/2025.01.23.634550

**Authors:** Lauren E Fries, Gabriel Grullon, Lauren Wilkes, Hanna E Berk-Rauch, Aravinda Chakravarti, Sumantra Chatterjee

## Abstract

Hirschsprung disease (HSCR) is a congenital enteric neuropathy caused by disrupted development of enteric neural crest-derived cells (ENCDCs). Although pathogenic coding variants in RET account for many cases, the largest genetic contribution to HSCR risk arises from a common non-coding variant (rs2435357) within a SOX10-bound RET enhancer (MCS+9.7) that reduces RET gene expression in vivo and triggers expression changes in other ENS genes in the human fetal gut. However, the ENS cell types affected by this enhancer and the mechanisms by which these transcriptional changes lead to HSCR remain unknown. Here, we investigated the role of this enhancer by generating mice carrying a deletion of the orthologous Ret mcs+9.7 enhancer (Δmcs+9.7). Single-cell RNA sequencing of E14.5 embryonic gut demonstrated that enhancer deletion reduced Ret expression by 8% without altering ENS cell composition. However, reduced Ret expression was restricted to differentiating neurons and inhibitory motor neuron lineages, revealing cell type-specific enhancer activity. To determine the functional consequences of further reducing Ret dosage, we generated compound heterozygous mice carrying both the enhancer deletion and a Ret coding null allele (+/Δmcs+9.7;+/CFP). These mice exhibited additive reductions in Ret expression, altered Sox10 expression, dysregulation of cell-cycle and neuronal differentiation programs, and selective depletion of developing inhibitory motor neuron lineages. These findings establish a cell type-specific role for the mcs+9.7 enhancer in modulating *Ret* dosage and reveal how subtle enhancer perturbations alter neural subtype specification without overt hypoganglionosis, suggesting that HSCR arises from a cascade of cellular defects triggered by >50% loss of Ret function.

## Introduction

The enteric nervous system (ENS), the largest component of the peripheral nervous system, is a ganglionic network that innervates the entire gastrointestinal tract. In mammals, this network contains hundreds of millions of neurons, including more than 600 million in humans.(Furness 2012). This network is formed by the cranio-caudal proliferation, differentiation and migration of Enteric Neural Crest-derived Cells (ENCDCs) in the developing gut across weeks 4 to 8 in the human and embryonic day (E) 10.5 to E14.5 in the mouse embryo (Furness 2012). One of the key genes involved in this developmental process is the receptor tyrosine kinase *RET* (Natarajan et al. 2002; Uesaka et al. 2008), whose expression in ENCDCs promotes proliferation and cell survival, although the gene is more broadly expressed in other tissues (e.g., kidney, brain, dorsal root ganglia). Significantly, coding and non-coding *RET* deficiency mutations lead to isolated Hirschsprung disease (HSCR) in ∼50% of cases, an uncommon (∼1/2,500-5,000 live births) developmental disorder of the ENS characterized by the absence of enteric ganglia along variable lengths of the distal bowel (Alves et al. 2013; Tilghman et al. 2019). In the developing mouse gut, *Ret* deficiency leads to significant transcriptional changes affecting numerous transcription factors (TFs), signaling molecules, and specific transport and biosynthesis genes, resulting in aganglionosis (Heanue and Pachnis 2006; Chatterjee et al. 2019). Analogous genetic changes are observed in the *RET* deficient human fetal gut with HSCR-associated susceptibility alleles in multiple *RET* enhancers (Chatterjee et al. 2023).

The cell type-specific *Ret* loss-of-function (LoF) phenotype has primarily been studied in *Ret* null homozygote mice (Lasrado et al. 2017; Vincent et al. 2023) which demonstrate that complete *Ret* deficiency leads to precocious differentiation and reduction in the number of proliferating ENCDCs thereby reducing the pool of progenitors available to generate later-developing neuronal populations, including the inhibitory motor neuron lineage. (Vincent et al. 2023). *Ret* heterozygous mice do not exhibit aganglionosis in the developing gut nor have any other phenotype associated with *Ret* LoF (Kapoor et al. 2017; Vincent et al. 2023). However, *Ret* heterozygote LoF in combination with hypomorphic homozygote mutations in the epistatic endothelin type B receptor gene (*EDNRB*) causes variable aganglionosis (Carrasquillo et al. 2002; McCallion et al. 2003).

In the human, most HSCR patients have heterozygous *RET* variants, some of which are complete LoF but most are hypomorphic; additionally, most patients harbor both *RET* coding and enhancer variants while yet others carry rare and common variants in other ENS and gut mesenchymal genes that are members of a large *RET-EDNRB* gene regulatory network (GRN) (Tilghman et al. 2019). Thus, we hypothesize that the totality of cis and trans sequence variants in an individual that reduce *RET* gene expression to less than 50% of wild type levels lead to functional *RET* haploinsufficiency and aganglionosis.

To address this hypothesis, we generated mouse models of HSCR with both coding and regulatory *RET* variants. Specifically, we first generated a mouse model by targeting an intronic enhancer of *Ret* (mcs+9.7) whose human homolog is the SOX10-bound *RET* enhancer (MCS+9.7) that harbors a common (24% - 45%) allele that individually increases HSCR risk 4-fold (Emison et al. 2005; Emison et al. 2010). Other *RET* enhancers also harbor polymorphic HSCR-associated variants which can cumulatively increase risk by 10-fold or greater (Chatterjee et al. 2021). Functionally, the common risk allele at MCS+9.7 disrupts SOX10 binding and leads to loss of *RET* function both in neural crest derived neuroblastoma cell lines (Chatterjee et al. 2021) as well as in the human fetal gut (Chatterjee et al. 2023). In addition, we also generated +/Δmcs9.7; +/CFP compound heterozygote mice by crossing the enhancer disrupted mice with a well characterized *Ret* null heterozygote mouse (*Ret^+/CFP^*) (Uesaka et al. 2008; Vincent et al. 2023), to study the progressive cellular defects arising from the coding and non-coding *Ret* variants.

We used single-cell RNA sequencing and quantitative RNA in situ hybridization to determine how enhancer and coding variants influence Ret expression across individual ENS cell populations and to identify the cellular mechanisms by which regulatory variation contributes to HSCR pathogenesis.

Our studies highlight that the HSCR phenotype results from precise thresholding of *Ret* gene expression below 50% in specific cell types in specific regions of the developing gut. However, HSCR is not a classical haploinsufficiency since the mutant phenotype can arise from more than one mutation affecting the same target gene (*Ret*). These results provide a basis for understanding the genotype-phenotype correlation and HSCR penetrance for different *RET* and other gene mutations in HSCR.

## Results

### Deletion of a HSCR-associated Ret enhancer in mice leads to loss of Ret expression in vivo

The MCS+9.7 *RET* enhancer, harboring the HSCR-associated common variant rs2435357, is sequence conserved in mice and other vertebrates (Emison et al. 2005; Grice et al. 2005), and functions as a *RET* enhancer in mouse and zebrafish transgenic assays (Grice et al. 2005; Chatterjee et al. 2016). MCS+9.7 binds SOX10 and regulates the expression of *RET* in human cell lines and the human fetal GI tract (Chatterjee et al. 2021; Chatterjee et al. 2023). First, we assessed the ability of this enhancer to regulate cell-type specific *Ret* expression in the developing mouse gut by deleting a 183 bp segment of the conserved region (Chr6:118,164,102-118,163,917; mm10) centered on the Sox10 binding site within the first intron of *Ret* (**Figure 1A; Supplemental_Fig_S1.pdf**). We measured *Ret* expression in the developing mouse gut, including the foregut, caecum, and hindgut, at E12.5 and E14.5, stages characterized by active enteric neural crest-derived cell migration, proliferation, and early neuronal differentiation. Quantitative real-time PCR (qRT-PCR) was performed with *Ret* expression normalized to the pan-neuronal marker *Tubb3*, to account for potential differences in neuronal abundance among genotypes. Heterozygous enhancer-deleted mice (+/Δmcs+9.7) showed no measurable change in Ret expression at either developmental stage. In contrast, Δmcs+9.7/Δmcs+9.7 homozygotes exhibited a small but reproducible reduction in neuron-normalized *Ret* expression, retaining approximately 94% of wild type expression at both E12.5 and E14.5 (*P*<0.01 for both) (**Figure 1B**).

**Figure 1.**
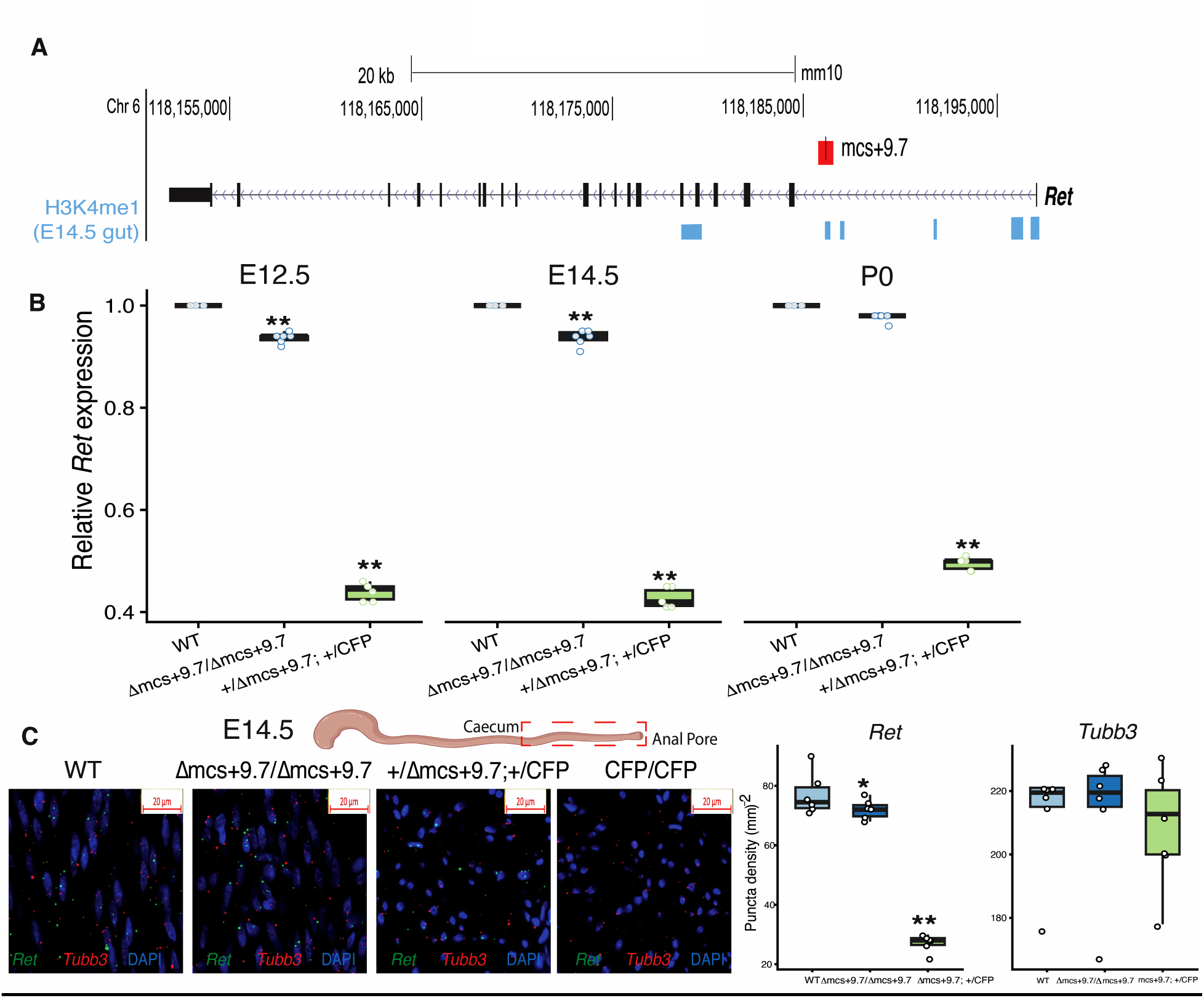
The mcs+9.7 enhancer quantitatively regulates *Ret* expression during embryonic ENS development. **(A)** Genomic organization of the mouse Ret locus (mm10 Chr6:118,155,000–118,195,000), showing the intronic mcs+9.7 enhancer and the 183-bp region deleted in the Δmcs+9.7 allele. H3K4me1 enrichment in E14.5 mouse gastrointestinal tissue is shown below the locus. **(B)** qRT-PCR measurement of Ret expression in the gastrointestinal tract at E12.5, E14.5, and P0. *Ret* expression was normalized to the pan-neuronal marker Tubb3 and expressed relative to the stage-matched wild type mean. Δmcs+9.7/Δmcs+9.7 embryos retained 94% of wild type *Ret* expression at E12.5 and E14.5 and returned to near-wild type levels at P0. Compound heterozygous +/Δmcs+9.7;+/CFP embryos retained 42% of wild type expression at E12.5 and E14.5 and approximately 50% at P0. Points represent biological replicates; boxes show the median and interquartile range. **(C)** Representative multiplex RNAscope images of E14.5 distal hindgut sections from wild type, Δmcs+9.7/Δmcs+9.7, +/Δmcs+9.7;+/CFP, and CFP/CFP embryos stained for *Ret* (green), *Tubb3* (red), and DAPI (blue). Box plots quantify *Ret* and *Tubb3* transcript punctas in wild type, Δmcs+9.7/Δmcs+9.7, and +/Δmcs+9.7;+/CFP embryos. Ret puncta density was progressively reduced across genotypes, whereas *Tubb3* puncta density was not significantly altered. \**P* < 0.05; \*\**P* < 0.01.

We next examined whether the contribution of the enhancer persisted at birth by measuring *Ret* expression at P0. At P0, *Ret* expression in Δmcs+9.7/Δmcs+9.7 mice increased to approximately 98% of wild type levels, indicating that the effect of mcs+9.7 deletion is largely restricted to embryonic ENS development. Thus, the 183-bp mcs+9.7 element functions as a quantitative developmental enhancer of *Ret* during the embryonic period when enteric progenitors are migrating, proliferating, and differentiating, but makes little detectable contribution to neuron-normalized *Ret* expression by birth.

To determine how enhancer loss interacts with reduced *Ret* coding dosage, we generated compound heterozygous mice carrying both the mcs+9.7 enhancer deletion and a *Ret* coding null allele. The coding null allele was generated by insertion of a Cyan Fluorescent Protein (CFP) cDNA into the first exon of *Ret* (*Ret^+/CFP^*) (**Methods** (Vincent et al. 2023)), producing +/Δmcs+9.7;+/CFP mice. These compound heterozygotes retained, on average, 42% of wild type Ret expression at E12.5 and E14.5, respectively, but increased to approximately 50% at P0 (*P*<0.01; Figure 1B). The embryonic expression levels were therefore lower than the approximately 50% expected from loss of one *Ret* coding allele alone, whereas the P0 level closely approached the expected heterozygous dosage. Because *Ret* expression was normalized to *Tubb3*, the embryonic reductions in Δmcs+9.7/Δmcs+9.7 and +/Δmcs+9.7;+/CFP mice cannot be attributed simply to a reduction in the number of enteric neurons. Instead, they indicate reduced *Ret* transcript abundance relative to neuronal content. Together, these findings demonstrate that mcs+9.7 contributes quantitatively to *Ret* transcription within the developing neuronal compartment and further reduces *Ret* expression when combined with a coding loss-of-function allele. The return of compound heterozygous expression to approximately 50% at P0 further supports the conclusion that the enhancer contribution is developmentally restricted and is largely absent by birth.

*Ret* expression is not restricted to ENS in the developing embryo but is expressed broadly in multiple neural and non-neural lineages during mouse embryogenesis. In situ hybridization and Northern blot analyses show that *Ret* transcripts first appear at embryonic day 8.5 and are detectable in a variety of developing tissues, including cranial ganglia, subsets of dorsal root and autonomic ganglia in the trunk, and motor neurons of the spinal cord and hindbrain, as well as undifferentiated neuroepithelial cells of the ventral neural tube and embryonic retina (Pachnis et al. 1993; Attie-Bitach et al. 1998). *Ret* expression is also observed outside the nervous system, most prominently in the excretory system including the nephric duct and ureteric bud epithelium during kidney organogenesis. To ascertain if the enhancer functions in the other non-GI tract *Ret* expressing cells, we also ascertained the tissue specificity of this enhancer by measuring gene expression of *Ret* in the developing kidney, an organ whose development is also dependent on the correct spatio-temporal expression of *Ret* (Schuchardt et al. 1994). There are no detectable changes in *Ret* expression in Δmcs+9.7/Δmcs+9.7; +/+ embryonic kidneys at either E12.5 or E14.5 but a 50% reduction in +/Δmcs+9.7;+/CFP mice, reflecting the loss of *Ret* gene expression from the heterozygous *Ret* knockout alone with no detectable additional contribution from the deleted enhancer (**Supplemental_Fig_S2.pdf**). These findings demonstrate that mcs+9.7 contributes to *Ret* expression in the developing ENS but has no detectable effect on *Ret* expression in the embryonic kidney at E12.5 or E14.5.

To determine whether the genotype dependent changes in *Ret* expression were detectable *in situ* within the developing ENS, we performed RNAscope for *Ret* and the pan-neuronal marker *Tubb3* in the E14.5 distal hindgut and quantified transcript puncta within matched regions of interest in the distal hindgut near the anal pore (see **methods**). *Ret* puncta density was modestly but significantly reduced in Δmcs+9.7/Δmcs+9.7 embryos compared with wild type controls, decreasing from a mean of approximately 77 in wild type to 62 puncta/mm². A substantially greater reduction was observed in +/Δmcs+9.7;+/CFP embryos, which retained only approximately 27 *Ret* puncta/mm² relative to wild type (*P*<0.01) (**Figure 1C**). In contrast, *Tubb3* puncta density was comparable among wild type, Δmcs+9.7/Δmcs+9.7, and +/Δmcs+9.7;+/CFP embryos, indicating that the reduced *Ret* signal was not explained by a corresponding reduction in neuronal transcript abundance within the analyzed regions. As expected, *Ret^CFP/CFP^*embryos showed an almost complete absence of *Ret* signal and a severe loss of enteric neurons in the distal hindgut, consistent with our previous findings (Vincent et al. 2023). To determine whether the reduced *Ret* signal simply reflected fewer enteric neurons, we normalized *Ret* transcript abundance to the neuronal marker *Tubb3* by calculating the *Ret/Tubb3* puncta ratio for each region of interest. The *Ret/Tubb3* ratio remained only modestly reduced in Δmcs+9.7/Δmcs+9.7 embryos and was markedly decreased in +/Δmcs+9.7;+/CFP embryos, confirming that the reduced *Ret* signal reflects diminished *Ret* transcript abundance within enteric neurons rather than differences in neuronal density (**Supplemental_Fig_S3.pdf**).

To determine if this cellular effect from reducing *Ret* expression is maintained at birth, we stained the descending colon, the derivate of the hindgut, with Acetylcholinesterase (AChE), the primary cholinergic enzyme found at postsynaptic neuromuscular junctions, and routinely used as a diagnostic marker for aganglionosis in humans (Yoshimaru et al. 2021) and the mouse (McCallion et al. 2003; Kapoor et al. 2017). We observed hypertrophic extrinsic fibers in the distal colon of +/Δmcs+9.7; +/CFP P0 mice reminiscent of short segment HSCR, with normal innervation observed in wild type and Δmcs+9.7/Δmcs+9.7 P0 colons (**Supplemental_Fig_S4.pdf**).

### Specific ENS transcriptional and cellular changes from functional loss of a Ret enhancer

We next explored which gut-specific cell type the enhancer was active in. To do so, we performed single-cell RNA-seq on dissected gastrointestinal tract (below the stomach to the anal pore) at E14.5, the developmental stage when ENS migration through the gut is complete and ENS cells have started differentiating into neuronal and glial subclasses. We studied tissues from each of two wild type and Δmcs+9.7/Δmcs+9.7; +/+ homozygote embryos to collect 25,028 wild type and 35,505 Δmcs+9.7/Δmcs+9.7; +/+ homozygote single cells, respectively (**Methods**). From these data we identified 3,415 high-variability genes and 20 principal components (PCs) to cluster cells based on their differential gene expression (*P*<0.05) across clusters relative to their mean expression in other clusters. Additionally, we used the following as marker genes to specify the identity of each cluster: Smooth muscle (*Acta2, Actg2, Myh11*; 32 % of cells), Epithelial (*Epcam, Cldn7, Lgals4*; 11%), Fibroblast (*Adamec1, Col6a4, Col3a1*; 30%), Endothelial (*Lars2, Tagln2*; 2%), Immune (*C1qa, C1qb, Cd52*; 2%), ENS (*Ret, Ednrb, Ngfr, Tubb3*; 11%) (Brugger et al. 2020; Fazilaty et al. 2021; Zhao et al. 2022) and a class of undifferentiated but transcriptionally active cells which have yet to acquire any major recognizable cell fate (*H9, Hmga2,Hoxa9*; 11%) (**Figure 2A; Supplemental_Fig_S5A.pdf**). Overall, there is no significant deviation in cell type composition between the wild type and the Δmcs+9.7/Δmcs+9.7 GI tracts (*P*=0.18; hypergeometric test) (**Figure 2B**). This is not surprising given that only 5% of Ret expression is lost in this genotype; note that in heterozygous Ret LoF mice (CFP/+) with 50% loss of Ret expression the phenotype is normal (Vincent et al. 2023).

**Figure 2.**
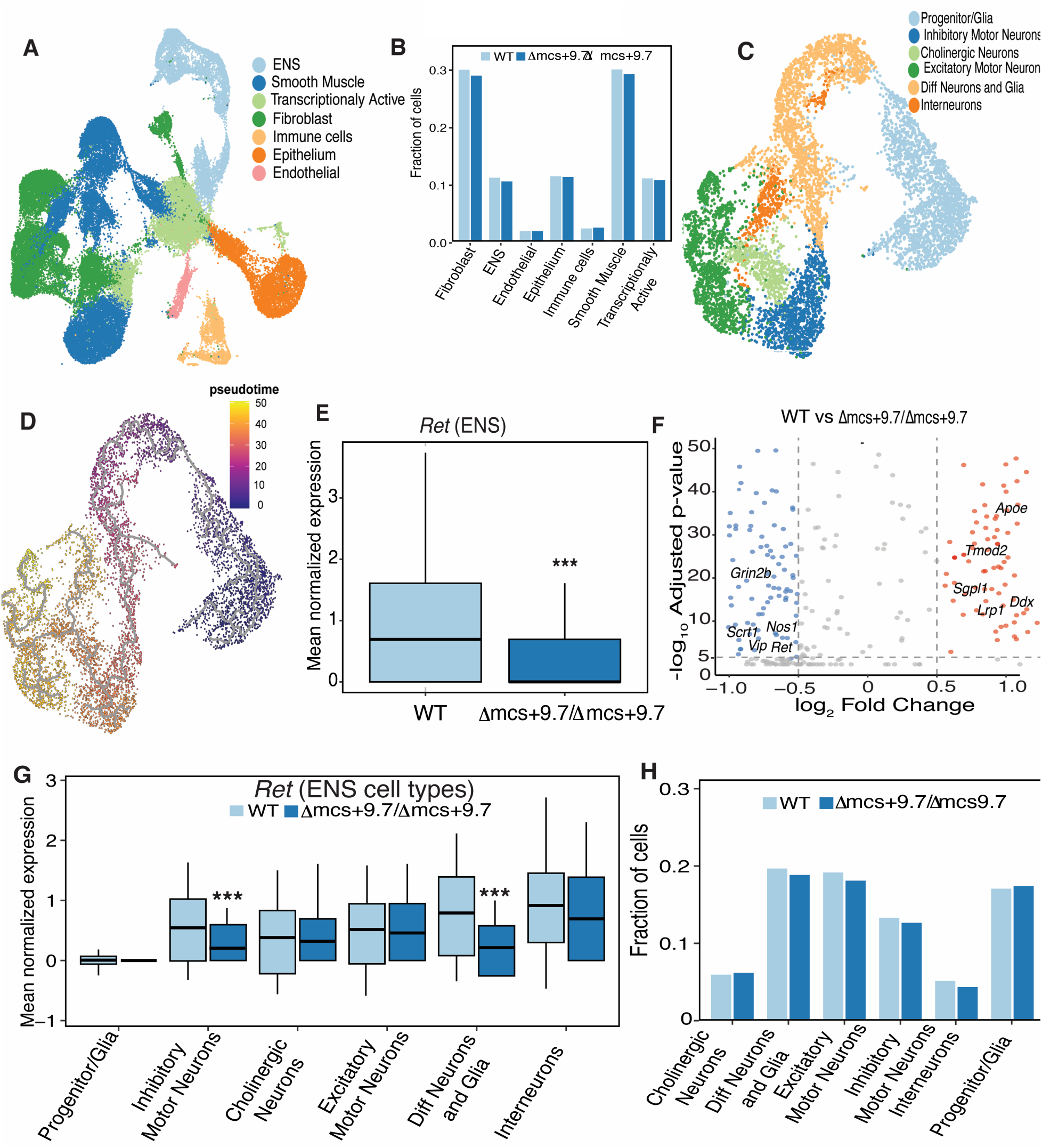
Homozygous deletion of mcs+9.7 alters ENS transcription without changing overall gut or ENS cellular composition. **(A)** UMAP representation of E14.5 gastrointestinal cells from wild type and Δmcs+9.7/Δmcs+9.7 embryos, colored by seven major cell populations. **(B)** Fractions of the major gastrointestinal cell populations in wild type and Δmcs+9.7/Δmcs+9.7 embryos; no significant global difference in composition was detected. **(C)** UMAP of the ENS compartment, colored by six annotated cell states. **(D)** Pseudotime analysis showing progression from progenitor/glial states toward differentiated neuronal states. **(E)** *Ret* expression across all ENS cells was modestly but significantly reduced in Δmcs+9.7/Δmcs+9.7 embryos relative to wild type**. (F)** Volcano plot of differential gene expression in the ENS of Δmcs+9.7/Δmcs+9.7 versus wild type embryos. Selected neuronal genes reduced after enhancer deletion and genes increased in the mutant are labeled. Dashed lines indicate the fold-change and adjusted-P-value thresholds used to define differential expression. **(G)** *Ret* expression within individual ENS cell states. Significant reductions were detected in inhibitory motor neurons and differentiating neurons and glia. **(H)** Fractions of ENS cell states in wild type and Δmcs+9.7/Δmcs+9.7 embryos; no significant depletion of any major ENS population was detected. Box plots show the median and interquartile range. Differential expression was assessed using the single-cell analysis described in Methods, with multiple-testing correction. \*\*\**P* < 0.001.

The ENS is the primary site of action for the *Ret* mcs+9.7 enhancer. The ENS cells we captured in our single cell experiments comprise 6 major cell types: (1) ENS progenitors with significantly higher expression (>2-fold; *P*<0.05) of canonical neural crest/progenitor markers (e.g*., Sox10, Ednrb*, and *Zeb2*) together with early glial-associated genes (e.g., *Plp1, Mpz*, and *Fabp7*). We have demonstrated these markers are co-expressed in embryonic enteric neural crest-derived progenitors before terminal glial differentiation, and therefore identify an immature progenitor population rather than mature enteric glia and combined use of these marker classes most accurately captures early ENS progenitors at embryonic stages, and avoids premature assignment to mature neuronal or glial identities (Vincent et al. 2023); (2) differentiating neurons and glia expressing pan neuronal and pan glial markers such as *Snap25, Plp1*, and *Mpz*, along with dorsal Hox transcription factors like *Hoxd5* and *Hoxd6*; (3) inhibitory motor neurons expressing *Nos1* and *Vip*; (4) excitatory motor neurons expressing *Calb2* and *Chat*; (5) interneurons expressing *Scrt1*, *Nxph4* and *Dlx5*; and, (6) cholinergic neurons expressing *Chat* and *Slc18a3* (**Figure 2C; Supplemental_Fig_S5B.pdf**). It is important to note that these neuronal populations represent developing embryonic neuronal lineages rather than fully mature neurons. Following established nomenclature in developmental ENS studies (Morarach et al. 2021; Vincent et al. 2023), we refer to these populations according to the neuronal subtype they are transcriptionally committed to generate (e.g., inhibitory motor neurons, excitatory motor neurons, cholinergic neurons), even though they remain immature and continue to undergo proliferation and differentiation at E14.5. These genes mark a developmental trajectory starting with progenitor cells that lead to more mature neuronal subtypes at the tip of the gene expression manifold with an intermediate state of actively dividing and differentiating neurons. To ascertain if the identified subclusters truly represent a developmental trajectory, we used their gene expression patterns to perform a formal trajectory analysis using Monocle 3 (Trapnell et al. 2014) which tests whether gene expression is associated with pseudo-time. This unsupervised reconstruction analysis does indicate that ENS cells follow the expected developmental path from undifferentiated (placed at time 0) to more mature neuronal cells (**Figure 2D**). Within the ENS cluster, *Ret* expression was reduced by 8% (*P*<0.001) (**Figure 2E**). Differential expression analysis of the ENS in wild type versus Δmcs+9.7/Δmcs+9.7 mice revealed 174 significantly affected genes (*P*<0.05), of which 51 were neuronal markers. These included the glutamatergic neuronal gene *Grin2b* (1.67-fold lower, *P*<0.01), interneuron markers *Scrt1* and *Nxph4* (1.28- and 1.37-fold reductions), and inhibitory motor neuron markers *Vip* and *Nos1* (0.6- and 0.8-fold lower, *P*<0.01 for all) (**Figure 2F**). Comparison with transcriptional profiles from complete *Ret* null mouse guts (Vincent et al. 2023) showed that 48 of these 174 genes overlapped with the 520 significantly affected genes (*P*<0.01) in *Ret*-expressing ENS cells, although the magnitude of change was smaller in the enhancer mutants, consistent with the milder allele (**Supplemental_Fig_S6.pdf; Supplemental_Table_S1.xlsx**). Notably, these transcriptional changes occurred without detectable alterations in ENS cell composition (**Figure 2B**), indicating that even a modest 5% reduction in *Ret* expression is sufficient to perturb a subset of the *Ret* associated transcriptional program observed in *Ret* null ENS cells, albeit with an attenuated response.

Conversely, 138 genes were significantly upregulated, enriched in two primary processes, regulation of cell size (*Tmod2, Lrp1, Shtn1, Cotl1*) and control of metabolic processes (*Sgpl1, Idh1, St8sia2*). Among these were also five genes with known roles in neurogenesis (*Mapt, Dcx, Dll3, Islr2, Apoe*). Together, these findings indicate that reduced *Ret* expression alters the ENS transcriptional landscape in a bidirectional manner, with downregulation of neuronal identity and function genes alongside upregulation of programs linked to growth, metabolism, and neurogenic or progenitor-associated states. Notably, these shifts are detectable at the transcriptomic level without an accompanying change in overall ENS cell composition at this stage, suggesting an early molecular perturbation that may precede, and potentially contribute to, later cellular or phenotypic consequences that are not yet apparent in our dataset.

Given *Ret*’s role in driving the differentiation of ENCDCs into more mature neuronal fates (Natarajan et al. 2002), our data show that *Ret* expression is nearly absent in progenitor cells at E14.5 in Δmcs+9.7/Δmcs+9.7 ENS but robustly expressed in all other cells (**Figure 2G**). Despite its broad expression in the developing ENS, in Δmcs+9.7/Δmcs+9.7, expression loss is limited to a 4-fold reduction (*P*<0.05) in differentiating neurons and glia and a 2.3-fold reduction (*P*<0.05) in inhibitory motor neurons compared to wild type (**Figure 2G**). These results reveal a cell type-specific regulation of *Ret* by the mcs+9.7 enhancer, confined to early differentiating cells and a specific neuronal subtype at E14.5. Importantly, although enhancer loss affects the transcriptional program in these cells, it does not lead to a detectable cellular phenotype, as identical proportions of cells were found in all clusters in both variant genotypes (**Figure 2H**).

### Coding and non-coding Ret variants lead to specific cellular defects in the ENS

A key question in HSCR is how lowering *Ret* gene expression levels below a threshold in the ENS lead to disease penetrance. It is clear that HSCR-associated variants within *RET* enhancers, and other enhancers at *NRG1* and *SEMA3* (Jiang et al. 2015), have lower penetrance than pathogenic *RET* coding variants, with the highest penetrance induced by *RET* null mutations (Tilghman et al. 2019). Indeed, disease segregation (penetrance) in some multiplex HSCR families can be better explained from the combined effects of coding and non-coding mutations at *RET* (Emison et al. 2010), reducing gene expression below a threshold. Therefore, we next assessed the effect of combined coding and non-coding *Ret* mutations (+/Δmcs+9.7; +/CFP) on the transcriptional and cellular architecture of the developing mouse gut. We dissociated cells from the gastrointestinal tracts of two E14.5 compound heterozygous embryos, generating 31,244 cells, of which 28,181 high-quality cells were retained after stringent quality control (Methods) and analyzed by UMAP using 2,000 highly variable genes and 20 principal components. These cells segregated into the same seven major gut cell types previously identified (**Figure 2A**), indicating that the overall cellular organization of the developing gut was preserved. However, unlike the Δmcs+9.7/Δmcs+9.7 embryos, the compound heterozygotes exhibited a measurable reduction in the proportion of ENS cells compared with both wild type and Δmcs+9.7/Δmcs+9.7 embryos (2% and 2.8% lower, respectively; *P* =0.001; hypergeometric test), whereas the remaining major gut cell populations were largely unchanged. This selective depletion of the ENS suggested that reducing *Ret* expression below the functional threshold primarily compromises the enteric lineage, prompting us to investigate the cellular and transcriptional consequences within the ENS in greater detail. We assessed 9,760 ENS cells in greater detail from all three genotypes, namely, wild type (n = 3,752 cells), Δmcs+9.7/Δmcs+9.7 (n = 3,183 cells), and +/Δmcs+9.7; +/CFP (n = 2,825 cells). These cells were re-clustered using 1,654 high variance genes and 20 PCs: we obtained eight clusters corresponding to the six major ENS cell types we had detected previously (**Figure 3A**). Combining cells from all 3 genotypes afforded us greater resolution to identify ENS cell subtypes since we could impute missing gene expression from the larger numbers of cells per cluster. The six clusters were annotated based on marker gene expression as Progenitor/Glia: (cluster 0: *Plp1, Mpz*; cluster 1: *Foxd3, Fabp7*; cluster 4: *Mest, S100b),* differentiating neuron and glia (cluster 2: *Cxcl12, Nkain4, Ednrb, Sox10*), inhibitory motor neurons (cluster 3: *Nos1, Vip*), cholinergic neurons (cluster 5: *Slc10a4, Chrnb2*), interneurons (Cluster 6: *Scrt1*, *Dlx5*, *Nxph4*) and excitatory motor neurons (Cluster 7: *Calb2, Chat*). Given our prior section’s observation that loss of the mcs9.7 enhancer selectively reduces *Ret* transcription in inhibitory motor neurons (iMNs) and differentiating neurons and glia, we thus asked whether combining enhancer loss with a heterozygous coding deletion (+/Δmcs9.7; +/CFP) would further deplete *Ret* expression in these specific embryonic cell types.

**Figure 3.**
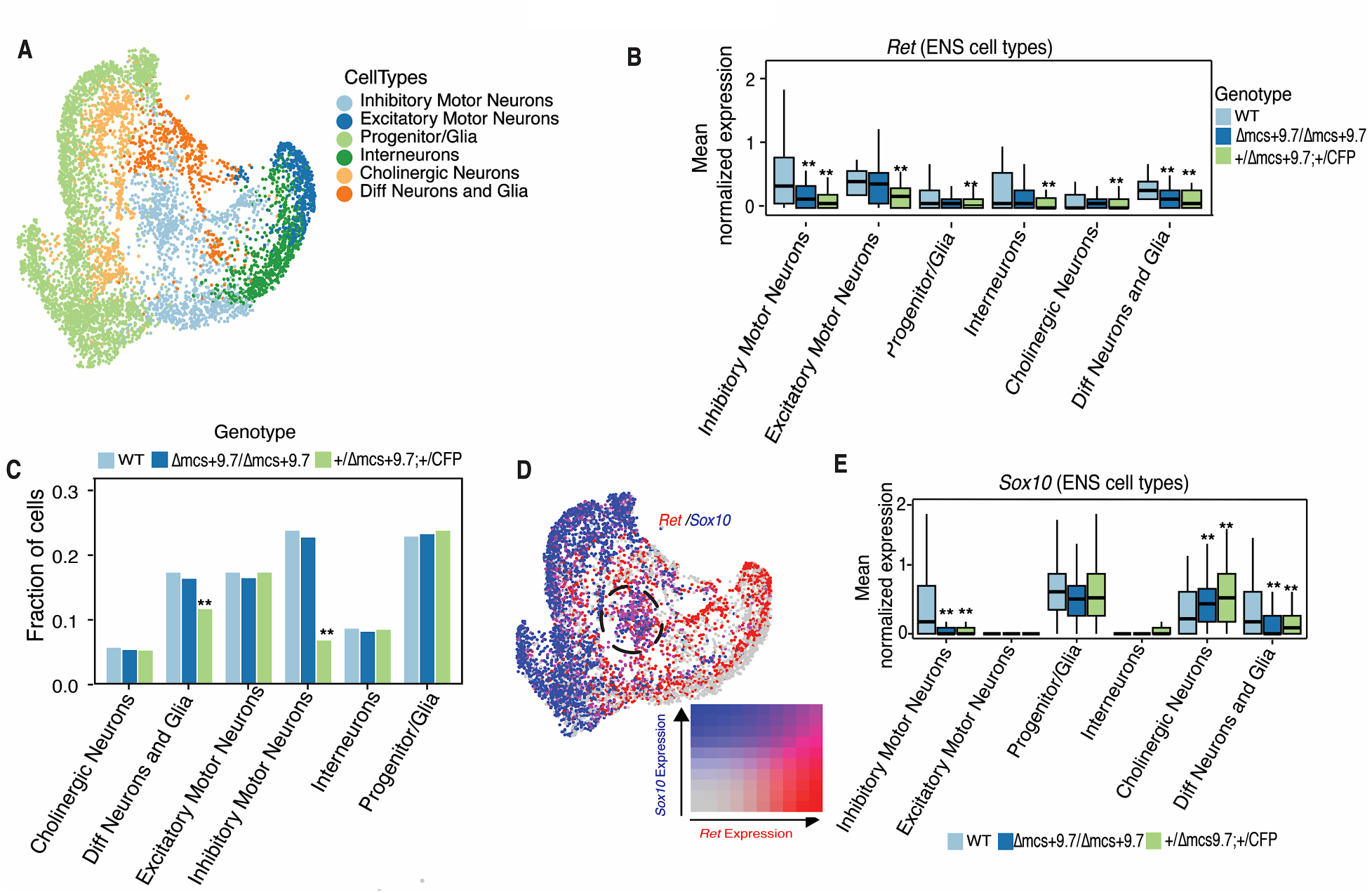
Combined enhancer and coding loss preferentially affects Ret-sensitive ENS lineages. **(A)** Integrated UMAP of E14.5 ENS cells from wild type, Δmcs+9.7/Δmcs+9.7, and +/Δmcs+9.7;+/CFP embryos, colored by six major ENS cell states. **(B)** Ret expression within each ENS cell state across the three genotypes. Compound heterozygous embryos showed the greatest reductions in inhibitory motor neurons and differentiating neurons and glia. **(C)** +/Δmcs+9.7;+/CFP embryos showed significant depletion of differentiating neurons and glia and inhibitory motor neurons (iMNs), while the remaining populations were comparatively preserved. **(D)** Bivariate UMAP visualization of Ret and Sox10 expression. Red indicates *Ret* expression, blue indicates *Sox10* expression, and purple indicates co-expression; the circled region highlights the iMN population, which contains the highest proportion of *Ret/Sox10*-coexpressing cells. **(E)** *Sox10* expression within each ENS cell state across genotypes. *Sox10* was reduced in inhibitory motor neurons and differentiating neurons and glia in compound heterozygotes, but increased in cholinergic neurons. Box plots show the median and interquartile range. Gene-expression comparisons were corrected for multiple testing; cell-proportion differences were assessed by chi-squared tests with P correction. \*\**P* < 0.01.

Indeed, in the compound +/Δmcs+9.7; +/CFP genotype, *Ret* expression was reduced well below the heterozygous coding baseline in a subset of ENS states, with a 3-fold decrease in inhibitory motor neurons (iMNs) and a 2.6-fold decrease in the differentiating neuron and glia population (*P* < 0.01 for both), demonstrating an additive effect of the non-coding enhancer and coding loss-of-function alleles (**Figure 3B**). In contrast, all other ENS cell types exhibited the expected ∼50% reduction in Ret expression (*P* < 0.01), consistent with heterozygous loss of the coding allele alone. Together, these data support a biphasic, cell-type-specific regulation of Ret by the mcs9.7 enhancer, wherein select ENS lineages display heightened sensitivity to combined regulatory and coding dosage perturbation. We then asked if decreasing *Ret* expression below 50% (below 0.5 -fold of wild type levels) had any effect on any ENS cell type. Using a chi-squared test for equality of cell proportions, we observed significant depletion of inhibitory motor neurons (39% lower: *P* = 3x10^-8^) and differentiating neurons and glia (34% lower: *P* = 2.2x10^-4^) in +/Δmcs+9.7 +/CFP embryos as compared to the wild type; all other clusters has the expected numbers of cells (progenitor cells: *P* = 0.4, excitatory neurons: *P* = 0.18, cholinergic neurons: *P* = 0.3, and interneurons: *P* = 0.3) (**Figure 3C)**. This cell depletion is observed in both embryos (replicates) increasing our confidence that this is not an artifact of cell sampling (**Supplemental_Fig_S7.pdf**). It is important to note that reducing *Ret* expression below 50% (approximately 0.5-fold of wild type) does not appear to affect the abundance of the early progenitor or glial population at this developmental stage. This observation is not unexpected, as we have previously shown that even complete loss of *Ret* (*Ret* null) does not consistently alter the size of this population up to E14.5 (Vincent et al. 2023). These findings suggest that changes affecting this compartment may occur later in development, beyond the window examined here. At the same time, we cannot fully exclude the possibility that subtle or highly specific subpopulations within the early progenitor or glial compartment are underrepresented or missed during cell capture across genotypes, which could mask more restricted effects of reduced *Ret* expression.

Given the increasing evidence pointing to the critical role of iMNs in ENS development (Morarach et al. 2021), and its specific loss attributed to aganglionosis in multiple animal models, including models of *Ret* deficiency (Spencer et al. 2021; Kuil et al. 2023; Vincent et al. 2023), we investigated if cellular loss of this cell type is a direct consequence of loss of *Ret* expression from its lowered enhancer activity. We calculated the proportions of cells which co-expressed both *Ret* and *Sox10*, the transcription factor bound to the mcs+9.7 enhancer, given Sox10’s role in regulating genes involved in maintaining progenitor/glial identity (Paratore et al. 2002; Bondurand and Sham 2013). This analysis was performed to identify ENS cell states most likely to be directly sensitive to Sox10-bound enhancer activity, and not to infer neuronal maturity or redefine cluster identity. *Ret/Sox10* co-expression likely reflects immature neuronal states, not mature neurons, and that this overlap is transient. Note that 88% of Sox10-expressing cells are in the progenitor or differentiating neurons and glia cluster (**Figure 3D**); conversely, 74% of *Ret*-expressing cells are specific lineage-committed neuronal cells of inhibitory (41%), excitatory (16%), cholinergic (12%) and interneurons (5%) (**Figure 3D**), highlighting *Ret*’s role in maintaining overall neuronal identity in the ENS (Natarajan et al. 2002). This analysis revealed, that in inhibitory motor neurons 83% of all cells co-express both genes, but only 18%, 2%, 0.4 % and 0.2% of differentiating neurons and glia, interneurons, excitatory motor neurons and cholinergic neurons, respectively, co-express *Ret* and *Sox10* (**Figure 3D**). We detected no cells co-expressing these genes in progenitor cells (**Figure 3D**). We further assessed *Sox10* expression to investigate potential transcriptional feedback between *Ret* and *Sox10*, as suggested by our earlier tissue-level gene expression studies in the gut. (Chatterjee et al. 2019). We observed a 1.8-fold and 1.6-fold reduction in Sox10 expression in iMNs and differentiating neurons and glia respectively in +/Δmcs9.7; +/CFP embryos compared to wild type (*P* < 0.01 for both), highlighting a cell type–specific positive feedback between *Ret* and its transcription factor *Sox10* (**Figure 3E**). Interestingly, we also detected a1.36 fold and 1.4-fold (*P*<0.01 for both) increased *Sox10* expression in cholinergic neurons in Δmcs9.7/Δmcs9.7 and +/Δmcs9.7; +/CFP respectively relative to wild type littermates (**Figure 3E**). This upregulation suggests the presence of a previously unrecognized, cell type–specific negative feedback mechanism. Together, these results indicate that the relationship between *Ret* and *Sox10* differs across ENS cell types. In inhibitory motor neurons and differentiating neurons and glia, reduced *Ret* expression is accompanied by reduced *Sox10* expression, consistent with positive regulatory coupling between the two genes. In contrast, *Sox10* expression increases in cholinergic neurons despite reduced *Ret*, suggesting that the interaction is different in this lineage. Thus, the effect of *Ret* deficiency on *Sox10* expression is cell type-specific rather than uniform across the ENS. Since reduced expression of *Ret* dysregulates feedback to *Sox10*, and both are critical drivers of aganglionosis in HSCR (Chatterjee et al. 2016; Chatterjee et al. 2019; Chatterjee et al. 2021), this suggest that iMNs and differentiating neurons and glia, which express both *Ret* and *Sox10*, are the primary cell types affected in HSCR.

Given that the phenotype of the compound heterozygotes could potentially be explained by heterozygous loss of the *Ret* coding allele alone, we next analyzed 2,813 ENS cells from E14.5 *Ret^+/CFP^* (*Ret* coding heterozygous) embryos together with the wild type, Δmcs+9.7/Δmcs+9.7, and +/Δmcs+9.7; +/CFP datasets (**Figure 4A**). Integration of the additional Ret+/CFP cells recovered all major ENS populations identified in the other genotypes, indicating that heterozygous coding loss does not eliminate any principal lineage at this developmental stage. The increased cell number also improved clustering resolution and revealed a small population of *Tubb3*-positive cells present across all four genotypes that lacked markers of inhibitory, excitatory, cholinergic, interneuron, progenitor, or glial identity. We therefore annotated this population conservatively as “neurons,” reflecting a neuronal state whose subtype identity could not be assigned with confidence from the available markers.

**Figure 4.**
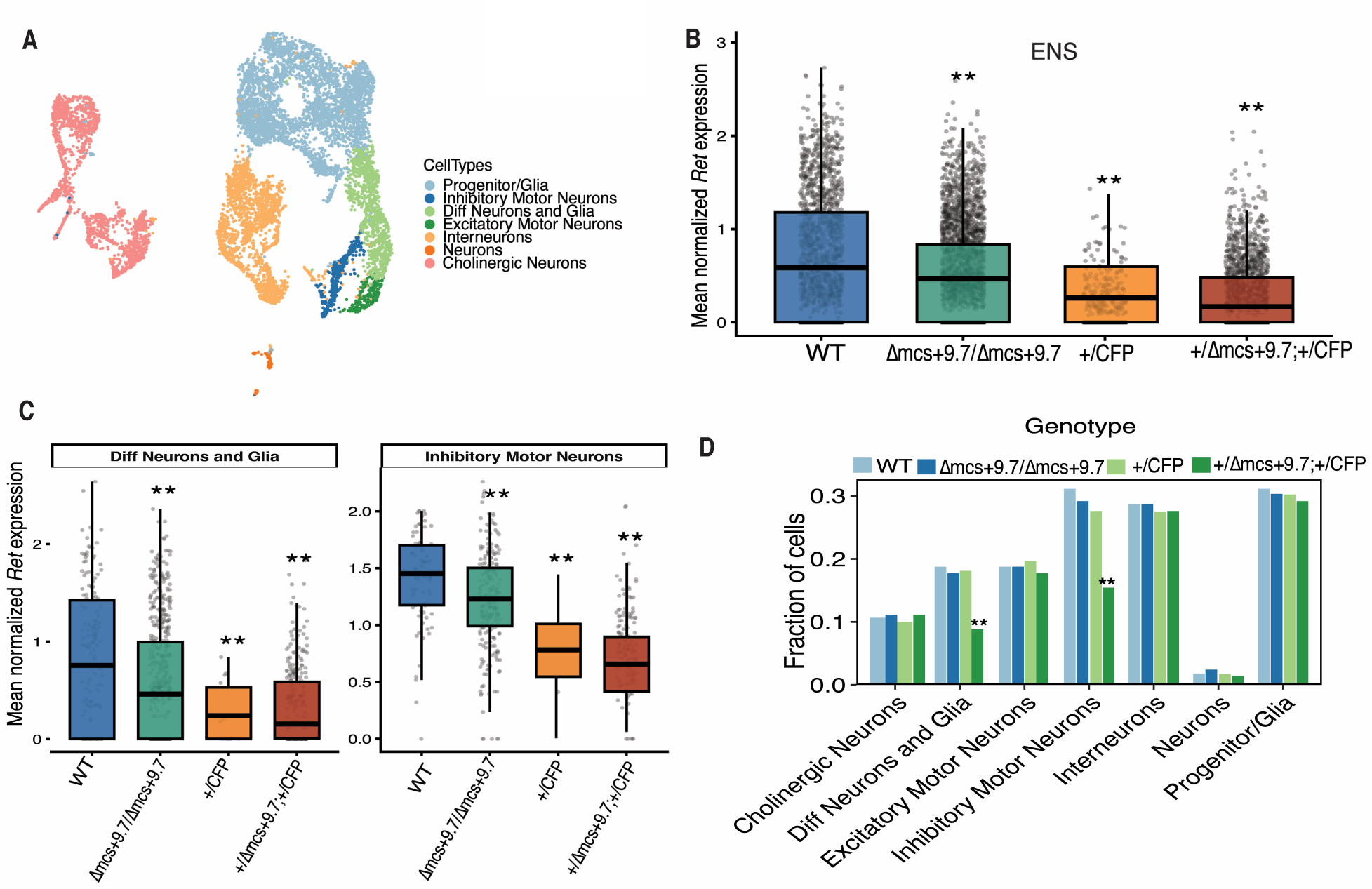
Ret coding heterozygosity alone does not reproduce the cellular phenotype of the compound genotype. **(A)** Integrated UMAP of E14.5 ENS cells from wild type, Δmcs+9.7/Δmcs+9.7, Ret+/CFP, and +/Δmcs+9.7;+/CFP embryos, colored by seven annotated ENS populations. **(B)** *Ret* expression across the complete ENS population. Expression decreased progressively from wild type to Δmcs+9.7/Δmcs+9.7, +/CFP, and +/Δmcs+9.7;+/CFP cells, with the compound genotype showing a further reduction beyond *Ret* coding heterozygosity alone. **(C)** The compound genotype showed the lowest expression in both enhancer-sensitive populations. **(D)** *Ret^+/CFP^* embryos retained near-wild type proportions of differentiating neurons and glia and showed no significant depletion of inhibitory motor neurons, whereas +/Δmcs+9.7;+/CFP embryos showed marked depletion of both populations. Box plots show the median and interquartile range; individual points in B and C represent cells. Gene-expression comparisons were corrected for multiple testing, and cell-proportion differences were assessed by chi-squared tests with P correction. \*\**P* < 0.01.

Across the complete ENS population, *Ret* expression decreased progressively with increasing genetic loss of Ret function. Relative to wild type, mean Ret expression across the entire ENS was reduced by approximately 50% in Ret+/CFP embryos (P<0.01) and by approximately 60% in +/Δmcs+9.7;+/CFP embryos (*P*<0.01), indicating that deletion of the mcs+9.7 enhancer produced an additional ∼10% reduction in Ret expression beyond coding heterozygosity alone (**Figure 4B**). This additional reduction was most apparent in inhibitory motor neurons, in which *Ret* expression was 48%(*P*<0.01) lower in +/CFP than in wild type cells and declined by a further 12% (*P*<0.01) in +/Δmcs+9.7;+/CFP. Differentiating neurons and glia also showed a marked reduction in *Ret* expression in both +/CFP (50% of wild type; *P*<0.01) and +/Δmcs+9.7;+/CFP embryos (42% of wild type; *P*<0.01), although the additional decrease in the +/Δmcs+9.7;+/CFP was more modest than that observed in inhibitory motor neurons (**Figure 4C**).

Importantly, *Ret* coding heterozygosity alone did not reproduce the cellular phenotype of the compound genotype. *Ret^+/CFP^*embryos retained approximately wild type proportions of differentiating neurons and glia and showed only non-significant reduction in inhibitory motor neurons. In contrast, +/Δmcs+9.7;+/CFP embryos exhibited a marked reduction in differentiating neurons and glia (approximately one-third lower than wild type and Ret+/CFP embryos), together with an approximately 50% depletion of inhibitory motor neurons (*P*<0.01). The proportions of the remaining ENS populations were comparatively preserved (**Figure 4D**). These results demonstrate that *Ret* heterozygosity alone is insufficient to explain the cellular defects observed in the compound mutants. Instead, the mcs+9.7 enhancer deletion further reduces *Ret* expression on the +/CFP background and pushes the two enhancer-sensitive ENS lineages below a functional Ret dosage threshold, resulting in their selective depletion.

To determine whether the additional reduction in *Ret* expression produced by the mcs+9.7 deletion was accompanied by broader molecular consequences, we compared the global ENS transcriptomes of wild type, +/CFP, and +/Δmcs+9.7; +/CFP embryos. Using an adjusted *P* value <0.05 and an absolute fold-change ≥1.5, we identified 590 differentially expressed genes (DEGs) between wild type and +/CFP ENS cells and 3,066 DEGs between wild type and +/Δmcs+9.7; +/CFP ENS cells. Importantly, direct comparison of +/CFP with +/Δmcs+9.7; +/CFP ENS identified 4,272 DEGs, including 1,417 upregulated and 2,855 downregulated genes, demonstrating that deletion of the mcs+9.7 enhancer produces widespread transcriptional changes beyond those caused by *Ret* coding heterozygosity alone. Comparison of the wild type and +/CFP, and wild type and +/Δmcs+9.7; +/CFP transcriptomes further revealed that 252 genes were differentially expressed in both genotypes, of which 82% changed in the same direction. These findings indicate that coding and regulatory reduction of *Ret* dosage perturb a common molecular program, while enhancer deletion further amplifies this transcriptional response on the +/CFP background. Together with the progressive reduction in *Ret* expression and selective depletion of differentiating neurons and inhibitory motor neurons, these findings support the mcs+9.7 enhancer as a quantitative modifier of Ret dosage that exacerbates, rather than redirects, the molecular consequences of Ret haploinsufficiency.

To define the biological processes underlying these transcriptional changes, we performed Gene Ontology enrichment analysis of the three pairwise comparisons. Despite differences in the individual DEGs identified in each comparison, all three datasets converged on a common set of neuronal developmental pathways, including axonogenesis, axon development, regulation of neuron projection development, cell morphogenesis involved in neuronal differentiation, synapse assembly, synapse organization, regulation of trans-synaptic signaling, modulation of chemical synaptic transmission, cell junction assembly, and regulation of cell adhesion (**Supplemental_Fig_S8.pdf**). These pathways were enriched both in the comparison of +/CFP and wild type ENS and, more significantly, following additional deletion of the mcs+9.7 enhancer on the +/CFP background, indicating progressive disruption of the molecular programs required for neuronal maturation and circuit assembly. Notably, direct comparison of +/Δmcs9.7; +/CFP and +/CFP ENS remained significantly enriched for many of these same processes, demonstrating that the enhancer contributes additional transcriptional perturbation beyond Ret coding heterozygosity alone. Collectively, these data indicate that the mcs+9.7 enhancer does not activate a distinct developmental program but quantitatively potentiates disruption of RET-dependent pathways governing enteric neuronal differentiation, axon extension, and synaptic maturation.

### Ret deficiency induces specific transcriptional changes in inhibitory motor neurons

Given the significant reduction in inhibitory motor neurons (iMNs) in +/Δmcs+9.7;+/CFP embryos, we next examined the transcriptional profile of this population independently from the remainder of the ENS (**Figure 5A**). These cells expressed the canonical inhibitory motor neuron markers *Nos1, Vip*, and *Gal*, while *Ntng1* marks a related neuronal population that arises developmentally from early nitrergic neurons (Morarach et al. 2021; Vincent et al. 2023) (**Figure 5B**). Differential expression analysis demonstrated that *Nos1, Vip*, and *Gal* expression was reduced by 2.8-, 1.6-, and 1.3-fold, respectively (*P*<0.01 for all comparisons), in +/Δmcs+9.7;+/CFP embryos, whereas *Ntng1* expression remained unchanged (Figure 5C). In contrast, Δmcs+9.7/Δmcs+9.7;+/+ embryos exhibited only modest reductions in *Vip* (0.6-fold) and *Nos1* (0.8-fold) expression (*P*<0.01 for both), with no significant changes detected in either *Gal* or *Ntng1* (**Figure 5C**). To determine whether these transcriptional changes identified by scRNA-seq were recapitulated *in vivo*, we performed RNAscope for *Nos1* and *Vip* on E14.5 distal hindgut sections from wild type, Δmcs+9.7/Δmcs+9.7, and +/Δmcs+9.7;+/CFP embryos and quantified transcript puncta within matched regions of interest (**Figure 5D**). Consistent with the single-cell analysis, Δmcs+9.7/Δmcs+9.7 embryos exhibited modest but significant reductions in both markers, with *Nos1* puncta density decreasing from approximately 123 to 102 puncta mm⁻² (17% reduction; *P*<0.01) and *Vip* puncta density decreasing from approximately 117 to 100 puncta mm⁻² (15% reduction; p<0.01) relative to wild type embryos. A substantially greater reduction was observed in +/Δmcs+9.7;+/CFP embryos, which retained only approximately 66 Nos1 puncta mm⁻² (46% reduction; *P*<0.01) and 49 Vip puncta mm⁻² (58% reduction; *P*<0.01) compared with wild type controls. These independent *in situ* measurements validate the transcriptional changes identified by scRNA-seq and demonstrate that progressive reduction of *Ret* dosage selectively compromises the inhibitory motor neuron gene program during ENS development.

**Figure 5.**
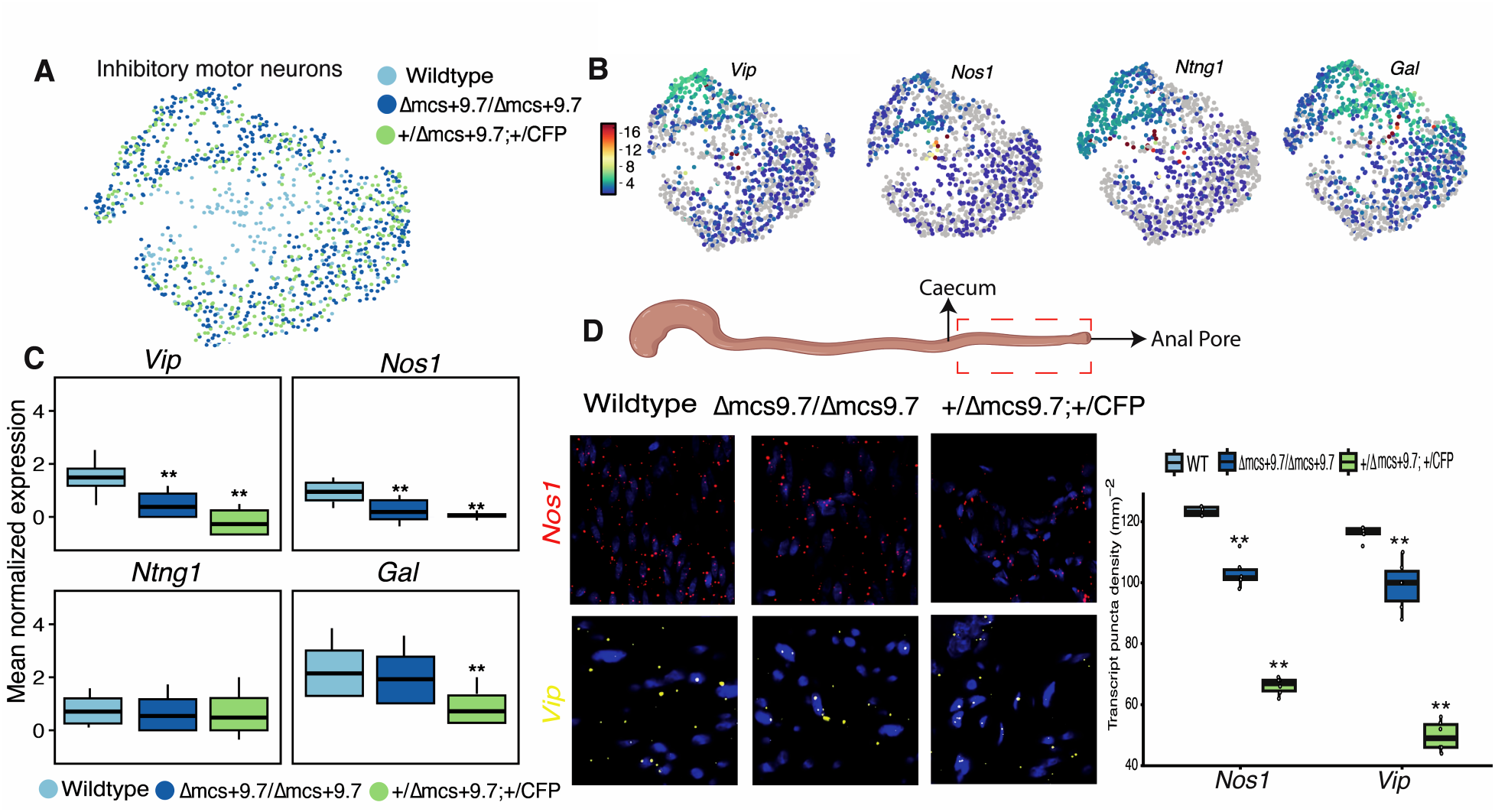
Progressive Ret deficiency disrupts the inhibitory motor neuron transcriptional program in vivo. **(A)** UMAP of E14.5 iMNs from wild type, Δmcs+9.7/Δmcs+9.7, and +/Δmcs+9.7;+/CFP embryos, colored bygenotype. **(B)** Feature plots showing expression of the inhibitory motor neuron markers *Vip, Nos1*, and *Gal* and the related nitrergic-lineage marker *Ntng1*. **(C)** Expression of *Vip, Nos1*, *Ntng1*, and *Gal* across genotypes. **(D)** Representative RNAscope images of E14.5 distal hindgut sections from the indicated genotypes stained for *Nos1* (red; upper row) or *Vip* (yellow; lower row), with DAPI counterstain (blue). Box plots show *Nos1* and *Vip* transcript puncta densities within matched regions of interest representing the median and interquartile range. RNAscope measurements were compared using Student’s t-tests. \*\**P* < 0.01.

### iMN specific effect in the HSCR specific GRN

Despite significant genetic heterogeneity, ∼67% of HSCR patients have mutations in genes of the *RET*-*EDNRB* gene regulatory network (GRN) (Tilghman et al. 2019). This GRN specifies bidirectional transcriptional feedback between *RET* and *EDNRB*, their transcription factors *SOX10, GATA2, RARB,* and *NKX2-5,* the *RET* co-receptor *GFRA1*, their ligands *GDNF* and *EDN3*, and the phosphorylated RET ubiquitin ligase degrader *CBL* (Chatterjee and Chakravarti 2019). To ascertain whether this feedback is associated with the iMN phenotype we observed, we quantified the expression of these genes and their changes in *Ret* deficiency. There is no expression of *Gdnf* and *Edn3* in iMN cells, as expected, since they are expressed and secreted from the surrounding mesenchyme (Baynash et al. 1994; Natarajan et al. 2002). *Ret* and its co-receptor *Gfra1* are downregulated 8-fold and 1.3-fold, respectively, while *Ednrb* is downregulated 6.2-fold (*P*<0.01 for all) in iMNs of +/Δmcs+9.7; +/CFP mice as compared to the wild type (**Figure 6A**). Only *Ret* is significantly (*P*<0.01) downregulated in the Δmcs+9.7/Δmcs+9.7 gut by 1.28-fold leaving both *Gfra1* and *Ednrb* unaffected. Among the transcription factors controlling *Ret* and *Ednrb*, we did not observe expression of *Gata2* and *Nkx2-5* in iMN cells at E14.5 but *Sox10* is reduced by 12-fold and 6-fold (*P*<0.01 for both), respectively, in +/Δmcs+9.7; +/CFP and Δmcs+9.7/Δmcs+9.7 iMN cells, respectively. Thus, both the modest 5% (0.95- fold) and the large 58% (0.42-fold) loss of *Ret* expression disrupt select components of the core *Ret-Ednrb* regulatory network in inhibitory motor neurons, reflecting heightened sensitivity of this lineage to Ret dosage reduction. Rather than indicating a global disruption of the HSCR gene regulatory network, these changes highlight lineage-restricted vulnerability of specific Ret-dependent regulatory nodes within iMNs has transcriptional changes affecting the same core GRN disrupted in HSCR leading to observable cellular changes in inhibitory motor neurons. In light of our results, it is unsurprising that the developmental trajectory leading to iMN formation is completely lost in *Ret* null embryos, as we have previously shown (Vincent et al. 2023).

**Figure 6.**
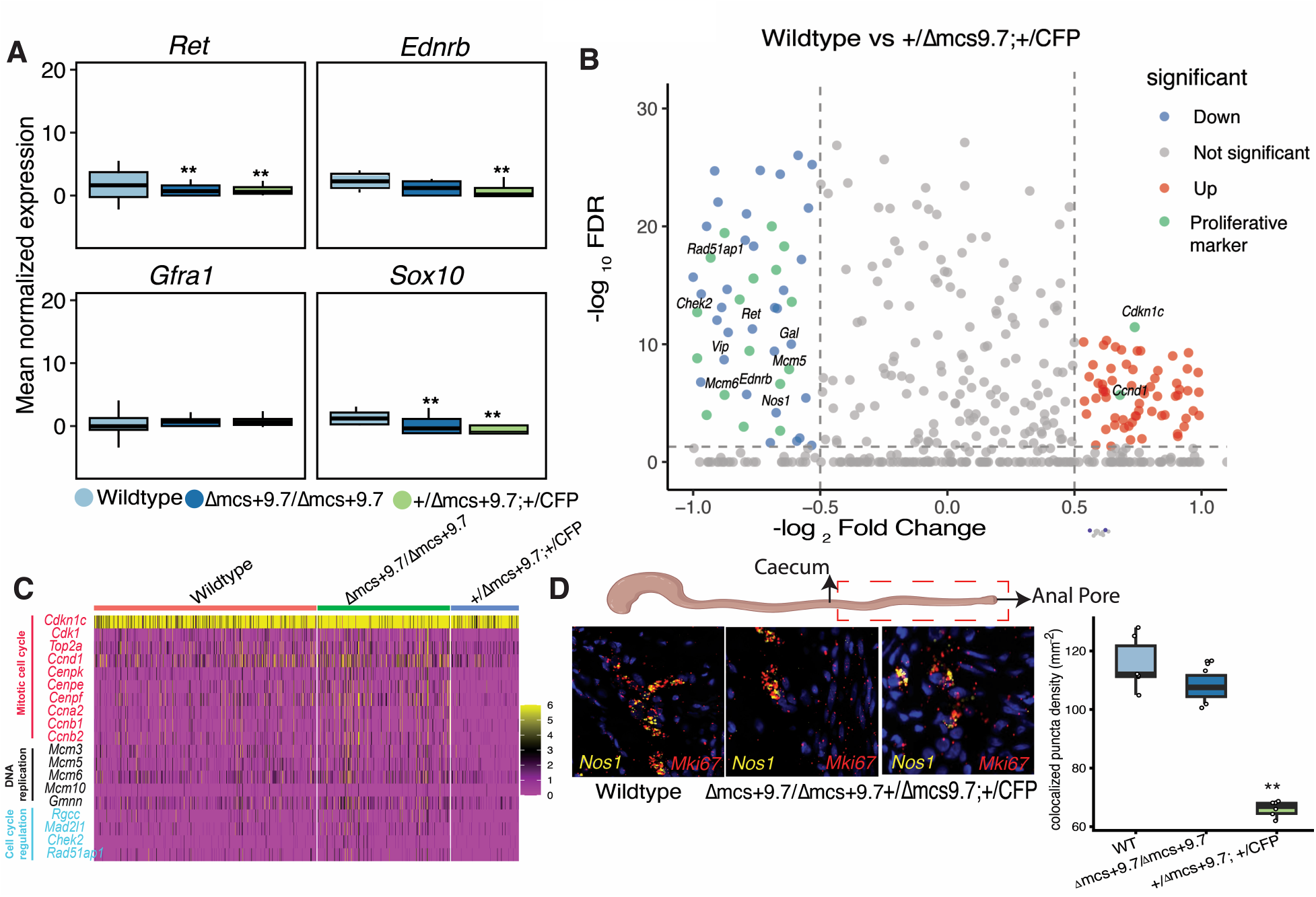
Reduced Ret dosage disrupts cell-cycle programs and proliferative capacity in inhibitory motor neurons. **(A)** Expression of *Ret, Ednrb, Gfra1*, and *Sox10* in inhibitory motor neurons from wild type, Δmcs+9.7/Δmcs+9.7, and +/Δmcs+9.7;+/CFP embryos. Ret was reduced in both mutant genotypes, whereas Ednrb, Gfra1, and Sox10 showed stronger disruption in the compound genotype. **(B)** Volcano plot of differential expression in inhibitory motor neurons from +/Δmcs+9.7;+/CFP versus wild type embryos. Downregulated genes are shown in blue, upregulated genes in red, nonsignificant genes in gray, and selected proliferation or cell-cycle associated genes in green. **(C)** Heat map showing expression of selected genes involved in mitotic cell-cycle progression, DNA replication, and cell-cycle regulation in inhibitory motor neurons from the three genotypes. **(D)** Representative multiplex RNAscope images of E14.5 distal hindgut stained for *Nos1* (yellow) and *Mki67* (red), with DAPI counterstain (blue). The diagram indicates the matched distal-hindgut region analyzed. The box plot shows the density of Nos1–Mki67 colocalized signals assigned to the same segmented cellular objects showing the median and interquartile range. RNAscope measurements were compared using Student’s t-tests. \*\**P* < 0.01.

### Ret deficiency decreases proliferative capacity in iMNs

Reduction in proliferative capacity resulting from complete loss-of-function of genes within the *Ret-Ednrb* gene regulatory network is a major driver of aganglionosis during ENS development (Bergeron et al. 2016; Fujiwara et al. 2018; Vincent et al. 2023). We therefore asked whether a similar, but quantitatively attenuated, mechanism underlies the selective loss of inhibitory motor neurons in +/Δmcs+9.7;+/CFP embryos. Differential expression analysis at E14.5 identified 93 genes (48 upregulated and 45 downregulated) that were significantly altered (*P*<0.05) in inhibitory motor neurons from +/Δmcs+9.7;+/CFP embryos compared with wild type littermates (**Figure 6B; Supplemental_Table_S2.xlsx**). These included ten genes involved in cell-cycle regulation, eight of which (*Ccnb2, Ccnb1, Ccna2, Cenpf, Cenpe, Cenpk, Top2a*, and *Cdk1*) were downregulated by >1.6-fold (P<0.01 for all), whereas *Cdkn1c* and *Ccnd1* were upregulated by 0.8- and 0.6-fold, respectively (P<0.01 for both) (Figure 6B). Genes involved in DNA replication (*Gmnn, Mcm3, Mcm6, Mcm5,* and *Mcm10*) and cell-cycle regulation (*Rgcc, Mad2l1, Chek2*, and *Rad51ap1*) were also downregulated by 1.4- to 2-fold (*P*<0.01 for all) (**Figure 6C and Supplemental_Table_S3.xlsx**). In Δmcs+9.7/Δmcs+9.7 embryos, all 19 genes changed in the same direction relative to wild type littermates, although not all reached statistical significance. Together, these data indicate that progressive reduction of *Ret* dosage disrupts both cell-cycle progression and neuronal differentiation programs within inhibitory motor neurons.

To determine whether these transcriptional changes were reflected *in vivo*, we performed RNAscope for the inhibitory motor neuron marker *Nos1* together with the proliferation marker *Mki67* (Seiler et al. 2019) in E14.5 distal hindgut and quantified colocalized transcript puncta within matched regions of interest (**Figure 6D**). Consistent with the transcriptomic data, Δmcs+9.7/Δmcs+9.7 embryos showed only a modest, non-significant reduction in *Nos1–Mki67* colocalization compared with wild type littermates (approximately 123 versus 107 colocalized puncta mm⁻², Student’s *t*-test, *P* = 0.24). In contrast, +/Δmcs+9.7;+/CFP embryos exhibited a marked reduction to approximately 66 colocalized puncta mm⁻², representing an approximately 46% decrease relative to wild type (*P* = 1.5 × 10⁻⁵) (**Figure 6D**). These independent *in situ* measurements validate the cell-cycle defects predicted by the single-cell transcriptomic analysis and demonstrate that reduced *Ret* dosage selectively impairs proliferation within the developing inhibitory motor neuron lineage.

To determine whether the reduced *Nos1* and *Vip* transcript abundance reflected a corresponding loss of inhibitory motor neurons, we quantified *Nos1⁺/Tubb3⁺* and *Vip⁺/Tubb3⁺* neurons from the same RNAscope datasets by assigning transcript puncta to individual segmented cells using *CellProfiler* (**Supplemental_Fig_S9.pdf**). Consistent with the transcriptomic analyses, Δmcs+9.7/Δmcs+9.7 embryos exhibited modest non significant reductions in both *Nos1⁺/Tubb3⁺* (1.5% decrease relative to wild type; Welch’s two-sample *t*-test, *P* = 0.052) and *Vip⁺/Tubb3⁺* neurons (1% decrease; *P* = 0.65). In contrast, +/Δmcs+9.7;+/CFP embryos showed substantially greater reductions, with *Nos1⁺/Tubb3⁺* and *Vip⁺/Tubb3⁺* neuronal densities decreased by 61.9% and 54.9%, respectively, compared with wild type littermates (*P* < 1 × 10⁻⁴ for both comparisons). These findings demonstrate that progressive reduction of *Ret* dosage results in a corresponding depletion of inhibitory motor neurons, confirming that the reduced *Nos1* and *Vip* transcript abundance reflects loss of these neuronal populations rather than simply decreased marker expression per neuron.

Collectively, these findings demonstrate that *Ret* dosage regulates ENS development through a transcriptional thresholding mechanism. Modest reductions in *Ret* expression, as observed in Δmcs+9.7/Δmcs+9.7 embryos, are sufficient to perturb neuronal gene expression programs but are largely buffered at the level of overall ENS cellular composition. However, further reduction of Ret dosage in +/Δmcs+9.7;+/CFP embryos amplify these same transcriptional perturbations beyond a critical threshold, resulting in selective defects in proliferation, differentiation, and ultimately cellular depletion. Importantly, this threshold is not uniform across the ENS. Although enhancer-null embryos show no detectable loss of total ENS cells, the earliest cellular consequences are already evident within the most Ret-sensitive neuronal lineage—the inhibitory motor neurons—where both molecular and subtle cellular changes occur in the same direction as those observed in the compound mutants. These data indicate that enhancer mediated modulation of *Ret* dosage first manifests as quantitative transcriptional changes, with overt cellular phenotypes emerging only after cell type-specific functional thresholds are exceeded.

## Discussion

Numerous genomic studies of complex disorders have implicated both rare coding and common noncoding variants as causal genetic factors for their etiology (Kapoor et al. 2014; Chatterjee et al. 2016; Kelly et al. 2022). However, it remains unclear how noncoding variants with small-to-modest genetic effects impact the same *phenotype* as their cognate large-effect coding variants. The latter are assumed to affect a rate-limiting gene’s function critically and therefore lead to a phenotype directly. But how do noncoding variants affect the same phenotype when the genomic evidence suggests that many such variants are required? We have previously hypothesized that this occurs only when the cumulative effect of all variants affect the same rate limiting step (a protein or a process modulated by several proteins) as the major coding mutations (Chakravarti and Turner 2016). We now provide evidence that, at least for HSCR, quantitative changes in *Ret* gene expression from both coding and noncoding variants, and its protein, affect the same gut developmental process in the same cells in the ENS at the same developmental time but do not affect all the target genes. The HSCR pathology (aganglionosis) results only when all target genes are affected as occurs through major qualitative changes in *Ret* gene expression and its protein.

The research described here provides some clues as to why these two genetic architectures, coding and noncoding, have different quantitative properties: higher vs lower penetrance; lower vs higher phenotypic variability. Despite affecting the same gut developmental program and the same GRN of genes, noncoding and coding *Ret* variants induce attenuated versus amplified responses of inhibitory motor neurons and a glial population with progenitor markers. Although such changes are now detectable at the single cell gene expression level, coding variants leads to phenotypic penetrance frequently while noncoding variants do so rarely. Why? A primary cause for this difference may be the stochastic nature of cell differentiation and proliferation, and *how many* cells are affected. Note that most but not all *RET* null heterozygotes have 100% penetrance (Emison et al. 2010). Thus, we believe that the number of cells which are affected, assumed to be those which have less than the optimum *Ret* gene expression of 50%, is larger in major coding mutations than in the common non-coding variants. Occasionally, the optimum is breached even with non-coding variants but this occurs rarely (lower penetrance).

Our results also demonstrate that the ‘aganglionosis’ detected in human patients, and currently tested for in mouse models, from acetylcholinesterase staining, might arise from loss of *specific cell types* and is not a broad loss of all neuronal cells, as previously thought (Furness 2012). This increases the stochasticity of penetrance because smaller numbers of cells are at risk increasing the role of functional ‘drift.’ Thus, pathophysiological studies of HSCR might benefit from the use of additional histopathological markers to assess the phenotype in patients. This comment pertains to both the aganglionic, hypoganglionic and transition zone segments of the gut.

One of the yet unsolved questions in HSCR is why do the lengths of aganglionic bowel (colon) differ in different patients, and why is short segment disease the most common (Badner et al. 1990)? Although heritable, many families have mixed short and long segment disease: the genetic basis of this segment length variation remains unclear but at least in some families arises from the interaction between *Ret* coding and enhancer variants (Emison et al. 2005; Emison et al. 2010). Previously studied mouse models have either total intestinal aganglionosis from complete *Ret* deficiency (Uesaka et al. 2008), which is extremely rare in humans, or variable long segment disease from multigenic models (McCallion et al. 2003; Hirst et al. 2017). Our double heterozygote mouse model with 42% (0.42-fold) *Ret* gene expression presents with aganglionosis only in the distal hindgut, thus most closely modeling short segment HSCR, the most common phenotype. An important consideration is that our analyses were intentionally focused on embryonic day 14.5, a developmental stage at which enteric neural crest cell colonization of the gut is essentially complete and neuronal subtype specification is actively underway. This developmental window allowed us to define the earliest cellular and transcriptional consequences of reduced *Ret* dosage before secondary pathological changes associated with aganglionosis become established. Although our findings demonstrate that the *Ret* mcs+9.7 enhancer functions during this critical period to regulate inhibitory motor neuron differentiation and ENS maturation, they do not exclude additional roles during later embryonic or postnatal development. Defining how these early molecular and cellular defects evolve over developmental time will be an important objective of future studies.

Finally, our data reaffirms previous observations that aganglionosis in HSCR animal models arises from proliferative defects in cells of the ENS. But why do all the cellular changes in these models affect only iMNs (Spencer et al. 2021; Kuil et al. 2023; Vincent et al. 2023)? One possible answer is that HSCR is basically a proliferative defect within developing neuronal precursors committed to the inhibitory motor neuron lineage and not a broad aganglionosis as previously assumed (Furness 2012). But suspicion remains that the human cellular phenotype may be more complex than in the mouse, since recent studies of ENCDCs derived from HSCR patient-induced pluripotent stem cells shows that both gene expression and cell type composition are different from mouse ENCDCs, with both committed glial progenitors (GP) and the neuronal progenitors (NP) absent in patient-derived ENCDCs; this feature has not been recapitulated in mouse models (Li et al. 2023). Given the limited sample size and technical variations in deriving ENCDCs this observed difference needs to yet be conclusively proven. To investigate these contrasting hypotheses directly, mouse models with human *RET* coding and noncoding normal and disease-associated haplotypes are indispensable.

More broadly, our findings illustrate the unique power of genetically engineered mouse models for dissecting the developmental consequences of human disease-associated coding and regulatory variants. Unlike human genetic studies, which identify statistical associations, or in vitro systems, which cannot fully recapitulate the spatiotemporal complexity of ENS development, mouse models enable direct investigation of gene dosage, enhancer function, cellular differentiation, and tissue morphogenesis within the developing embryo. By combining precise coding and enhancer alleles *in vivo*, we demonstrate that common regulatory variants can quantitatively modify the effects of rare coding mutations on the same developmental program, thereby providing a mechanistic explanation for the variable penetrance and expressivity characteristic of Hirschsprung disease. This framework extends beyond *RET* and HSCR and provides a general strategy for understanding how coding and noncoding variation interact during mammalian development to shape the phenotypic diversity of complex congenital disorders.

## Materials and Methods

### Nomenclature of mutant mice

Our mouse models include disruptions in both the coding and enhancer regions of the same gene; an approach not commonly used in human genetic studies. To ensure clarity, as well as correspondence between the human and mouse orthologous elements, we provide the nomenclature table below, which we have consistently applied throughout the manuscript.

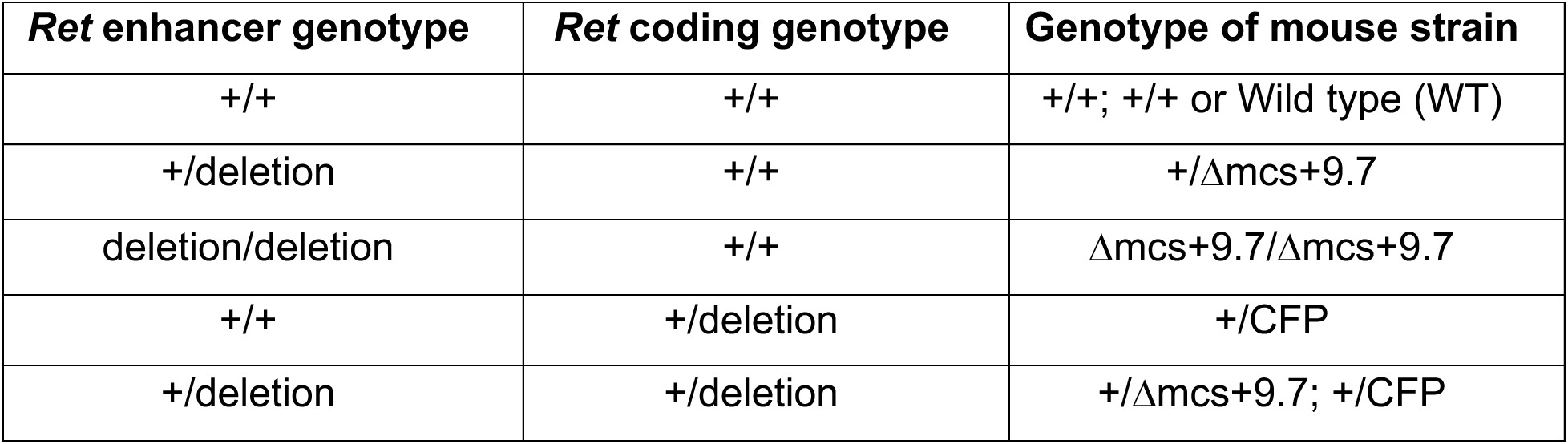

### Generation of Ret deficient mouse models

#### mcs+9.7 deletion mice

We used the following guide RNAs to delete the human *RET* enhancer MCS+9.7 (Chr10: 43,581,812–43,582,711; hg19) conserved in mice (mcs+9.7: 6:118,164,102-118,163,917; mmu39) with sequence identity of 88%: Forward Guide: TCCTTCCGGCTTCTAGACAA (Chr 6:118163913 – 118163932) and Reverse Guide: GCAGCAGCAACAGCTACTTG (Chr 6:118164067 – 118164086). The guides were designed to delete a DNA fragment within the conserved sequence centered on the known Sox10 binding site. They were injected with Cas9 protein into one cell embryos by standard methods at the NYU transgenic core facility to delete the target DNA segment. We used the following primers to routinely genotype these mice: mcs+9.7 mKO-F: CACGTCCTGTCTCTTCCGTAG and mcs+9.7 mKO-R: AGCCTGTGAACGTGTTCTGC. Both the heterozygous and homozygous mcs+9.7 deletion (Δ) mice were born in the expected Mendelian proportions (**Supplemental_Table_S4-8.xlsx**). These are referred to as Δmcs+9.7/+; +/+ and Δmcs+9.7/Δmcs+9.7; +/+, respectively.

#### *Ret* CFP knock-in mice (*Ret^CFP/+^;* MGI:3777556)

These mice were generated by crossing *Ret*fl/+ mice to β-actin-Cre mice to remove the RET9 cDNA: these mice were maintained on a C57BL/6 background and have been previously described in detail (Vincent et al. 2023). To generate double heterozygote mice deleted for both the mcs+9.7 enhancer and the coding sequence of *Ret*, +/Δmcs+9.7; +/+ mice were crossed to +/+; *CFP/+* mice to produce Δmcs+9.7/Δmcs+9.7; +/+ and +/Δmcs+9.7; +/CFP genotypes (the mutant sites are listed as mcs+9.7 and coding, respectively); all offspring were in expected Mendelian proportions (**Supplemental_Table_S4-8.xlsx**). All mouse experiments were conducted in accordance with the NIH Guidance for the Care and Use of Laboratory Animals. All procedures were approved by the NYU School of Medicine Animal Care and Use Committee (Protocol number: s17-01779).

### Dissection and dissociation of embryonic tissue

Foregut and hindgut tissue from wild type, Δmcs+9.7/Δmcs+9.7, +/CFP and Δmcs+9.7/Δmcs+9.7; +/CFP embryos at E14.5 were dissociated into single-cell suspensions using Accumax (Sigma, USA). The cells were filtered serially through a 100 μm and 40 μm cell strainer, and centrifuged at 2,000 rpm for 5 m. The cell pellets were resuspended in 5% FBS, 4 mM EDTA in Leibovitz L-15 medium. This cell suspension was diluted to 20,000 cells and processed through the 10x Genomics GEM generator to create a standard 3’ library using the 10x Genomics standard protocol. These cells were sequenced at an average of 1x10^5^ ± 10^4^ reads/cell.

### Single-cell gene expression (RNA-seq) analysis

Raw sequencing data were processed using the 10x Genomics Cell Ranger pipeline to generate cell barcode, feature, and gene-expression count matrices. These matrices were imported into *Seurat* (Stuart et al. 2019) or downstream quality control, normalization, dimensionality reduction, clustering, and comparative analysis. Each genotype was initially processed independently. Cells were retained if they satisfied the following quality-control criteria: ≥500 unique molecular identifiers, ≥250 detected genes, log10 genes per UMI >0.80, and mitochondrial transcript fraction <20%. Genes detected in fewer than 10 cells were excluded from downstream analyses. Gene-expression data were normalized and variance stabilized using SCTransform in Seurat (Hafemeister and Satija 2019) which also identified highly variable genes for dimensionality reduction and clustering. To minimize technical and batch-associated differences while preserving biologically shared and genotype-specific cell states, datasets from the individual genotypes were integrated using Seurat’s anchor-based integration framework. This procedure first identifies pairs of transcriptionally similar cells across datasets, termed integration anchors, and then uses these anchors to align shared cellular states across experimental conditions. The integrated expression space was used for principal-component analysis, graph-based clustering, and uniform manifold approximation and projection to generate a two-dimensional representation of the cellular landscape.

Cell clusters were annotated on the basis of established lineage and subtype marker genes. Comparative analyses of cell-type abundance and gene expression were performed across genotypes within the integrated dataset. Unless otherwise indicated, plotted gene-expression values represent the average normalized transcript abundance within the specified cell population, calculated from the normalized single-cell expression matrix. These values therefore reflect the magnitude of expression across cells within a population and not merely the proportion of cells in which Ret transcripts were detected. All raw and processed single cell data files are available at Gene Expression Omnibus (GEO) under accession number GSE304252. The analysis pipeline used for *Seurat* can be found at https://github.com/SumantraChatt/mouse-scRNAseq/tree/main.

### Single-cell differential gene expression analysis

We used Seurat’s *FindMarker* feature to perform differential expression by pseudo-bulking the samples either across genotypes or across cell clusters. We utilized “MAST”: GLM-framework (Finak et al. 2015) that treated the cellular detection rate as a covariate within *Seurat*.

### Cell-type annotation

Cell identities were assigned using a combination of unsupervised differential expression analysis, functional enrichment, and established lineage markers. Marker genes for each cluster were identified using Seurat by comparing each cluster against all remaining cells. Genes exhibiting >1.5-fold differential expression at a false discovery rate (P) <0.05 were considered cluster-enriched markers. Functional annotation of these marker sets was first performed using DAVID(Huang et al. 2007) and significantly enriched Gene Ontology (GO) biological processes (P <0.01 containing >10 genes) were used to infer the major biological identity of each cluster. As an independent validation, GO and Kyoto Encyclopedia of Genes and Genomes (KEGG) pathway enrichment analyses were performed using clusterProfiler (v4.16.0)(Yu et al. 2012) to confirm the functional annotation of each population. For enteric nervous system (ENS) populations, cluster identities were further refined using established lineage- and subtype-specific marker genes rather than relying solely on pathway enrichment. Annotation incorporated canonical markers for ENS progenitors, differentiating neurons and glia, excitatory and inhibitory motor neurons, interneurons, cholinergic neurons, and enteric glia, based on published single-cell atlases of the developing ENS (Elmentaite et al. 2021; Morarach et al. 2021; Vincent et al. 2023). Final annotations therefore integrated unbiased transcriptomic signatures, functional pathway enrichment, and prior biological knowledge to assign robust cell identities.

### RNAscope fluorescent in situ hybridization and image acquisition

Whole gastrointestinal tracts extending from immediately distal to the stomach through the terminal hindgut were dissected from E14.5 embryos and fixed in 4% paraformaldehyde. Tissues were processed for paraffin embedding, sectioned at 10-µm thickness, and mounted on positively charged glass slides. Multiplex fluorescent RNA in situ hybridization was performed using the RNAscope Multiplex Fluorescent Reagent Kit v2 and mouse-specific target probes from Advanced Cell Diagnostics, according to the manufacturer’s protocol for formalin-fixed, paraffin-embedded tissue. Briefly, sections were deparaffinized, treated with hydrogen peroxide, subjected to heat-mediated target retrieval and protease digestion, and hybridized with probe sets targeting *Ret* (Catalog # 431798; Bio-Techche), *Tubb3* (Catalog # 423398), *Nos1(*Catalog # 437658), *Vip (*Catalog # 415968), or *Mki67 (*Catalog # 416778), as appropriate for each experiment. Probe signals were sequentially amplified and detected using horseradish peroxidase-mediated tyramide signal amplification, after which nuclei were counterstained with DAPI and slides were mounted using an antifade mounting medium.

Images were acquired using a a Nikon Eclipse Ti confocal microscope and processed using Fiji. Confocal stacks were taken using a, 40X oil objective with Type A immersion under identical acquisition settings for all samples within each probe combination. For image presentation, stacks were z-projected to single images using the maximum value for each pixel confocal microscope with a 40× objective Analyses were restricted to anatomically matched regions of interest in the distal hindgut near the prospective anal pore. Multiple nonoverlapping fields were acquired from each embryo.. The number of embryos and regions of interest analyzed for each experiment are indicated in the corresponding figure legends.

### Image Quantification

A custom CellProfiler pipeline, adapted from a previously described RNAscope analysis workflow (Erben et al. 2018), was used to quantify transcript puncta. Multichannel images were separated into their constituent fluorescence channels, and DAPI-positive nuclei were identified using the ‘*IdentifyPrimaryObjects*’ module with global Otsu thresholding. Nuclear objects were filtered on the basis of size, shape, and fluorescence intensity to exclude debris and nonspecific background. Because no membrane marker was included, approximate cellular territories were generated by isotropic expansion of individual nuclei and defined using the ‘*IdentifySecondaryObjects’* module.

Signals corresponding to *Ret, Tubb3, Nos1, Vip*, and *Mki67* were independently enhanced, segmented, and filtered using channel-specific thresholds optimized for punctum size, intensity, and local background. Detected puncta were assigned to their corresponding nuclear or approximated cellular parent objects using the ‘*RelateObjects*’ module. Channel-specific correction was applied when necessary to minimize autofluorescence, background signal, and fluorescence bleed-through. Transcript counts, object intensities, spatial coordinates, and parent–child relationships were exported for downstream analysis.

For measurements of transcript density, the total number of puncta detected within each region of interest was divided by the analyzed tissue area and reported as puncta per mm². *Ret* abundance was additionally normalized to neuronal transcript content by calculating the *Ret/Tubb3* puncta ratio within each matched region of interest. For analysis of the developing inhibitory motor neuron lineage, *Nos1* and *Vip* puncta densities were quantified independently. Colocalization of *Nos1* and *Mki67* was assessed by identifying puncta assigned to the same approximated cellular object and reporting the density of *Nos1–Mki67*-positive objects or colocalized signals per mm². Quantified measurements were imported into R for statistical analysis and visualization using ggplot2. Statistical significance was assessed using a paired Student’s t-test.

### Gene expression TaqMan assays

Total RNA was isolated from whole gut tubes at E12.5 and E14.5 and from the distal colon at P0 using TRIzol reagent (Thermo Fisher Scientific) and purified using the RNeasy Mini Kit (QIAGEN) according to the manufacturers’ instructions. For kidney expression analyses, both kidneys were dissected from each embryo at E12.5 and E14.5, pooled, and processed similarly for RNA isolation. Five hundred nanograms of total RNA from each sample were reverse transcribed using SuperScript IV Reverse Transcriptase (Thermo Fisher Scientific) with Oligo(dT) primers. cDNA was diluted 1:5 and analyzed by TaqMan quantitative PCR using gene-specific assays for *Ret* (Mm00436305_m1; Thermo Fisher Scientific). To normalize for differences in enteric neuronal abundance between genotypes, *Ret* expression was quantified relative to the pan-neuronal marker *Tubb3* (Mm01250302_g1; Thermo Fisher Scientific). Relative expression was calculated using the ΔΔCt method and expressed relative to stage-matched wild type controls. Five independent biological samples were analyzed for each genotype and developmental stage, with each biological sample assayed in technical triplicate. Statistical analyses were performed using the biological replicates (n = 5).

The mean of the ΔC_t_ values (C_t_ Condition - C_t_ Actin) for each replicate for each condition was calculated after converting them to a linear scale (2^Δct^). The fold change was calculated as the ratio of the mean of 2^Δct^ experiment / 2^Δct^control. The mean for the control was set to unity and the relative fold change was calculated based on these values. For calculating the variance of the of the fold change we used the following formula: 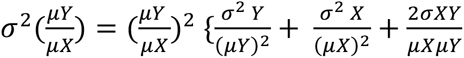

where *σ*^2^ is variance, *μ* is mean and 2*σXY* is the covariance between an experimental condition Y and its corresponding control X. Subsequently, *P* values were calculated from pairwise 2-tailed t-test, and the data presented as the fold change with its standard error.

## Data Access

All raw and processed single cell data files are available at Gene Expression Omnibus (GEO) under accession number GSE304252.

## Acknowledgement

This work was supported by NIH grants DK135089 to AC and SC, HD116004 to SC, and HD028088 to AC. The funders had no role in design of the study or data interpretation. SC and AC conceived and designed the study. LEF and HRB created all mouse constructs. LEF, GG, LW conducted single-cell sequencing and RNA in situ assays; SC and LEF analyzed the data. SC and AC wrote the manuscript. All authors approved the final version of the manuscript.

## Declaration of Interests

The authors declare no competing interests.

## Notes

### Competing Interest Statement

The authors have declared no competing interest.

### Summary of Updates

We have performed extensive new experiments and analyses, including single-cell RNA sequencing of the ENS from Ret heterozygous (Ret+/CFP) embryos and quantitative RNAscope in situ hybridization to independently validate the major gene expression changes identified by our transcriptomic analyses.

