## Supplemental Figures for "Combined effects of *Ret* coding and enhancer loss-of-function alleles cause progressive loss of inhibitory motor neurons in the enteric nervous system"

### Supplemental Figure 1

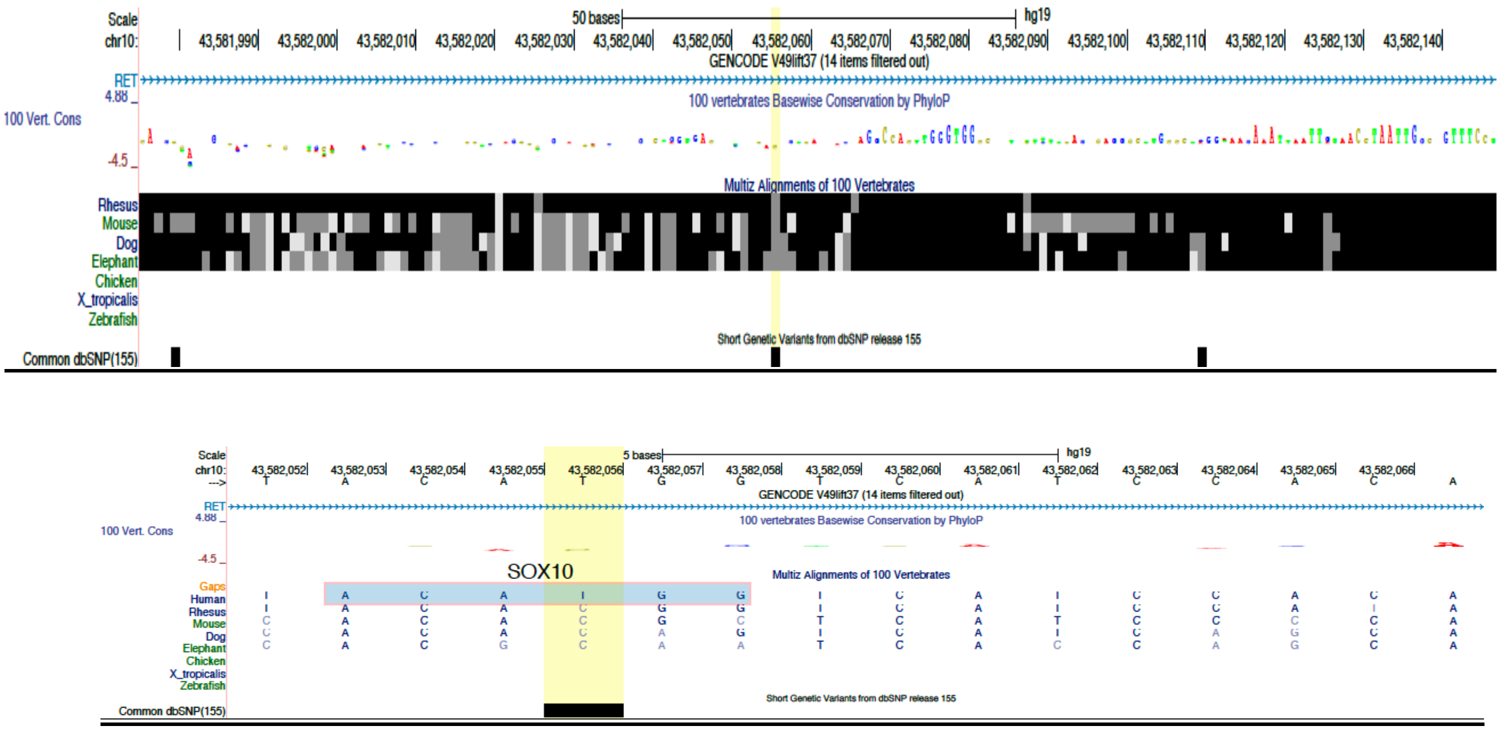


**Supplemental Figure 1**: The human genomic interval chr10:43,581,976–43,582,147 (GRCh37), containing the *RET* intronic enhancer mcs9.7, shows high sequence conservation across vertebrate species, with 80 percent conservation based on PhyloP scores and multiz alignment. The vertical yellow line marks the position of the Hirschsprung disease associated polymorphism rs2435357. A magnified view of the region is shown below, highlighting a conserved SOX10 transcription factor binding site. The HSCR risk allele, "*T*", disrupts this binding motif, consistent with reduced RET enhancer activity observed in disease associated haplotypes.

#### Supplemental Figure 2

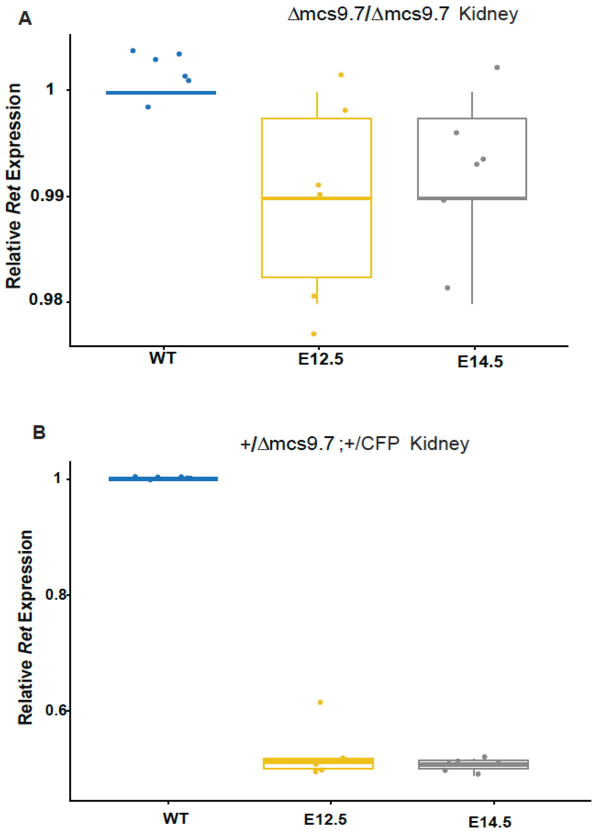


**Supplemental Figure 2:** (A) qPCR analysis Ret gene expression in developing mouse kidneys at E12.5 and E14.5 in the Δmcs+9.7/Δmcs+9.7 embryos show no significant change in *Ret* expression. (B) Similar analysis in the +/Δmcs+9.7;+/CFP embryos at same developmental stages show 50% reduction in Ret expression accounted for by the Ret null allele (CFP) but effect from the enhancer.

**Supplemental Figure 3**

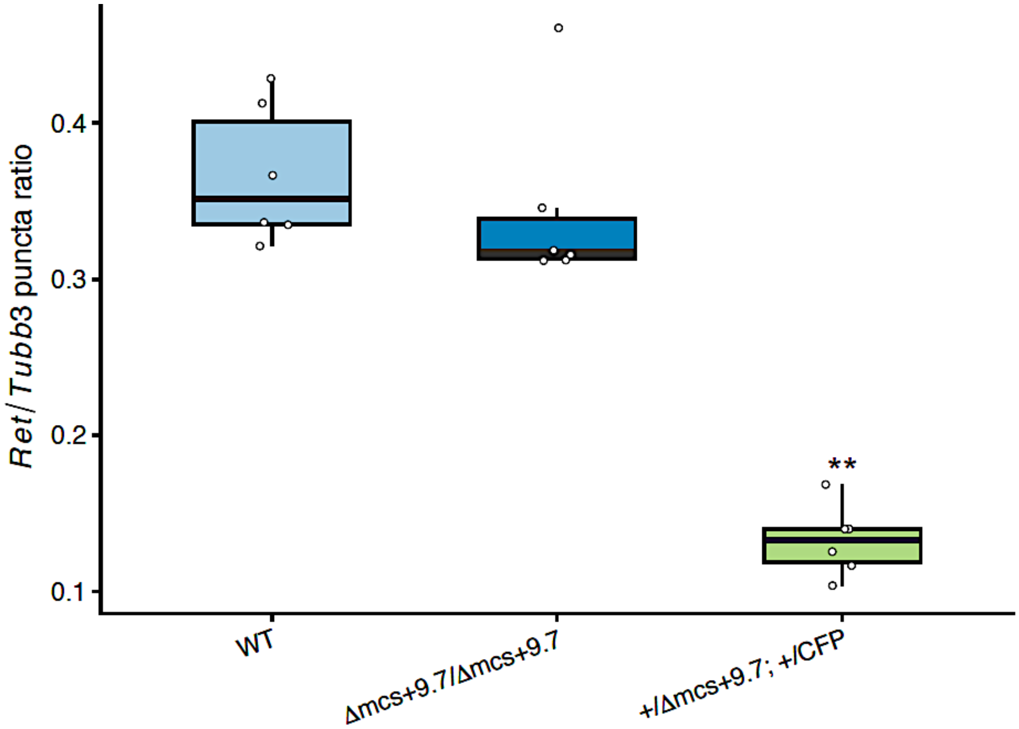


**Supplemental Figure 3:** Quantification of the *Ret/Tubb3* transcript puncta ratio in the E14.5 distal hindgut determined by RNAscope. *Ret* transcript abundance was normalized to the pan-neuronal marker *Tubb3* for each quantified region of interest. The modest reduction in Δmcs+9.7/Δmcs+9.7 embryos and the marked decrease in +/Δmcs+9.7;+/CFP embryos confirm that reduced *Ret* expression is not solely explained by differences in neuronal abundance. Boxes represent the median and interquartile range; each point represents an independently quantified region of interest. P values were calculated relative to wildtype using a two-tailed Student's *t*-test; P < 0.01 (**)

Supplemental Figure 4

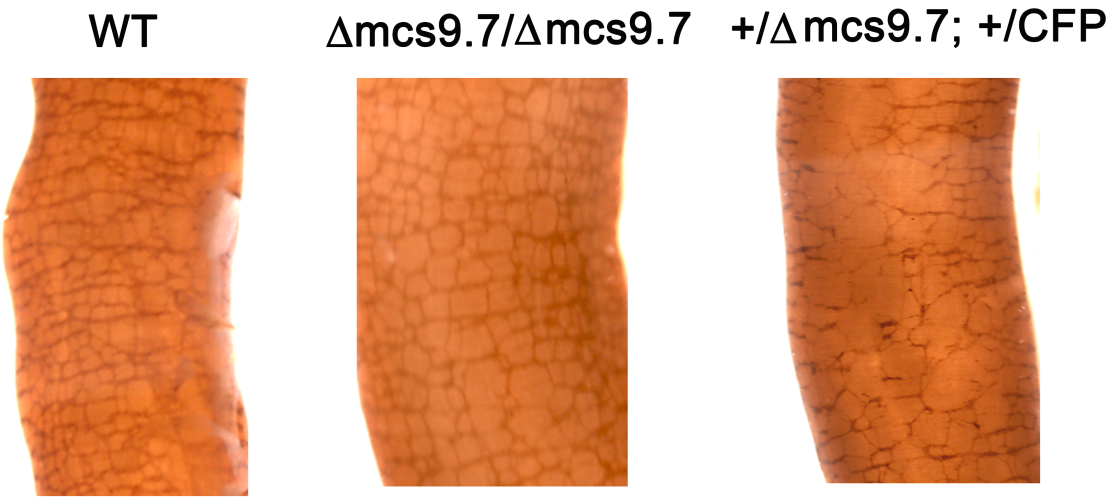


**Supplemental Figure 4:** Acetylcholinesterase (AChE) staining for cholinergic neuron fibers at postsynaptic neuromuscular junctions in mice at birth (P0) reveals significant reduction of cholinergic neuron fibers in the distal colon of +/Δmcs+9.7;+/CFP reminiscent of short segment Hirschsprung disease, with normal innervation observed in wildtype and Δmcs+9.7/Δmcs+9.7 mice.

#### Supplemental Figure 5

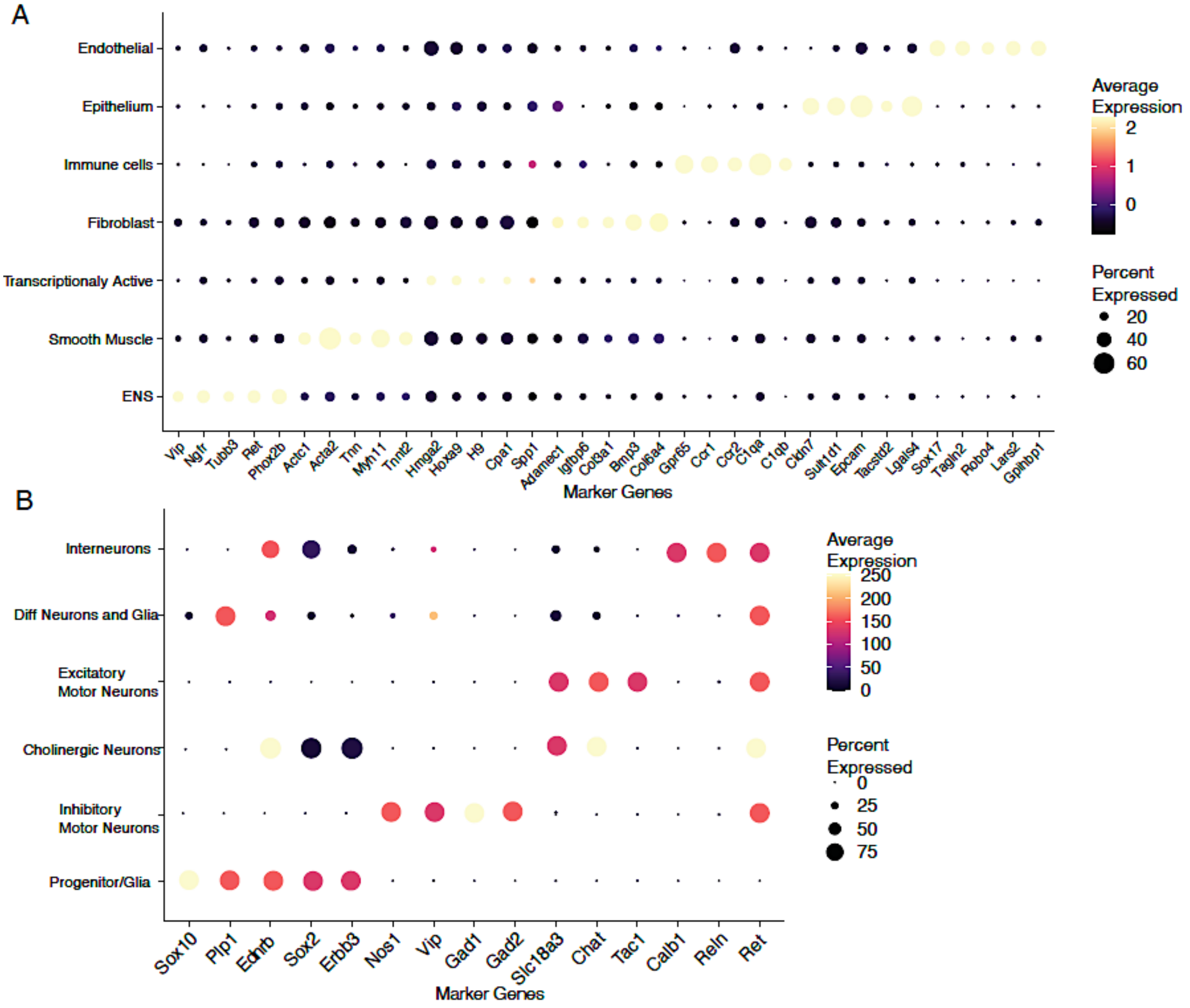


**Supplemental Figure 5:** (A) Dot plot showing representative marker genes used to annotate major cell populations in the E14.5 mouse gut, including enteric nervous system (ENS), smooth muscle, fibroblasts, immune, epithelial, and endothelial cells. (B) Dot plot showing marker genes distinguishing ENS subtypes, including progenitor/glial cells, inhibitory and excitatory motor neurons, cholinergic neurons, interneurons, and differentiating neurons and glia. In both panels, dot color indicates average gene expression, and dot size represents the percentage of cells within each cluster expressing the gene.

**Supplemental Figure 6**

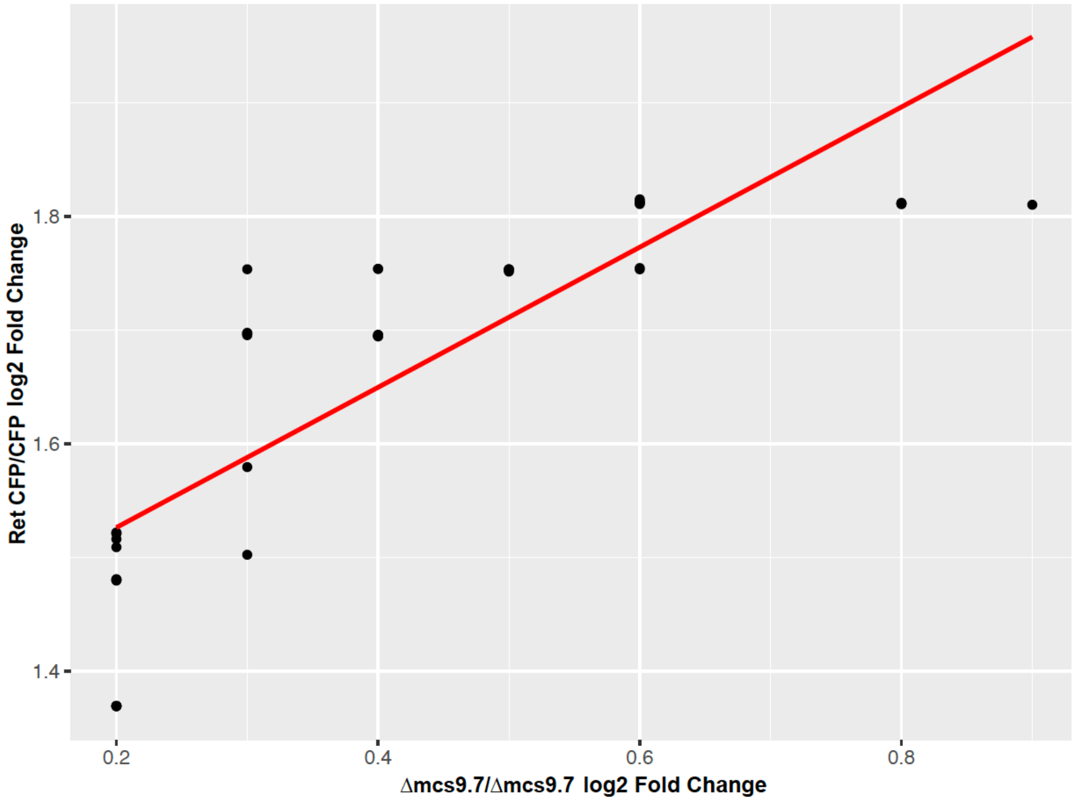


**Supplemental Figure 6:** 48 genes ENS expressed genes show similar direction of gene expression changes in both the Δmcs+9.7/Δmcs+9.7 mice (95% Ret expression) and the $Ret^{CFP/CFP}$ (0% Ret expression)) albeit with much attenuated effect.

#### Supplemental Figure 7

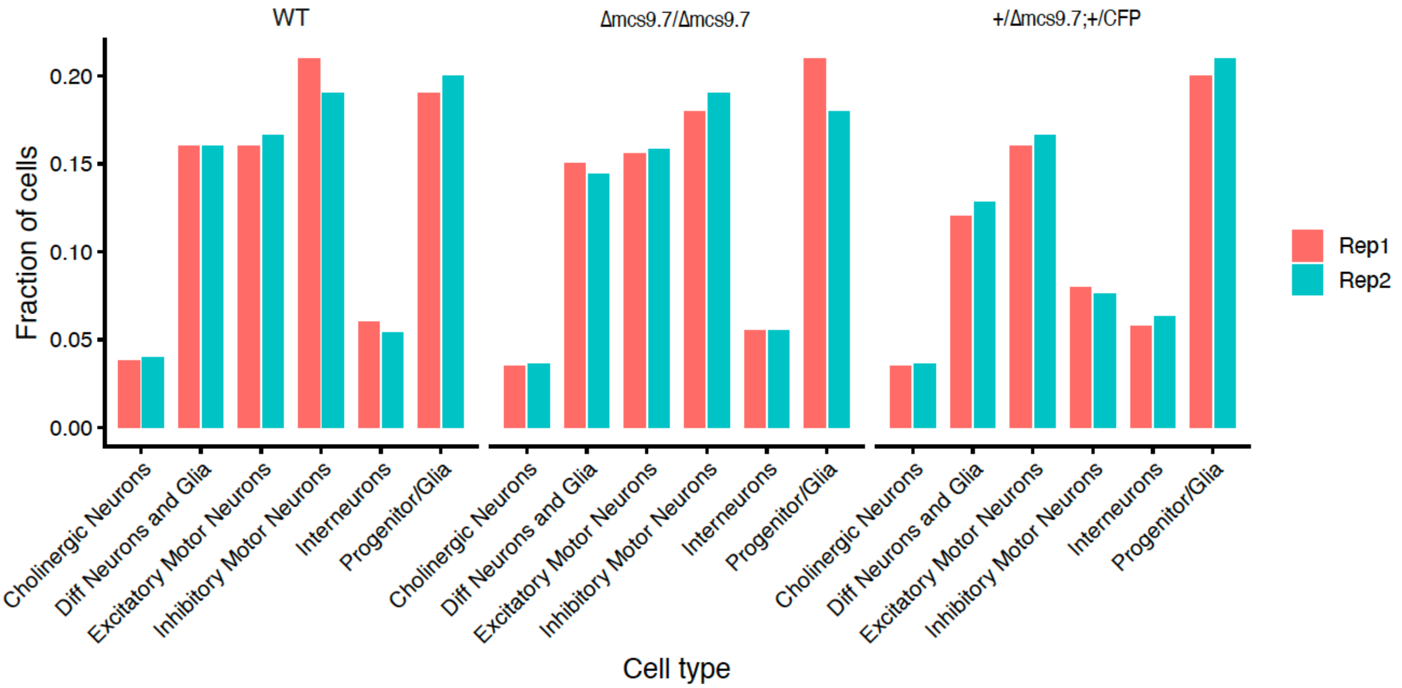


**Supplemental Figure 7:** Bar plots show the fraction of each enteric nervous system (ENS) cell type detected in individual biological replicates for wildtype (WT), Δmcs9.7/Δmcs9.7, and compound heterozygous (+/Δmcs9.7; +/CFP,) embryos at E14.5. Each bar represents a single embryo (Rep1 or Rep2), plotted separately to explicitly illustrate replicate-level consistency rather than averaged values.

#### Supplemental Figure 8

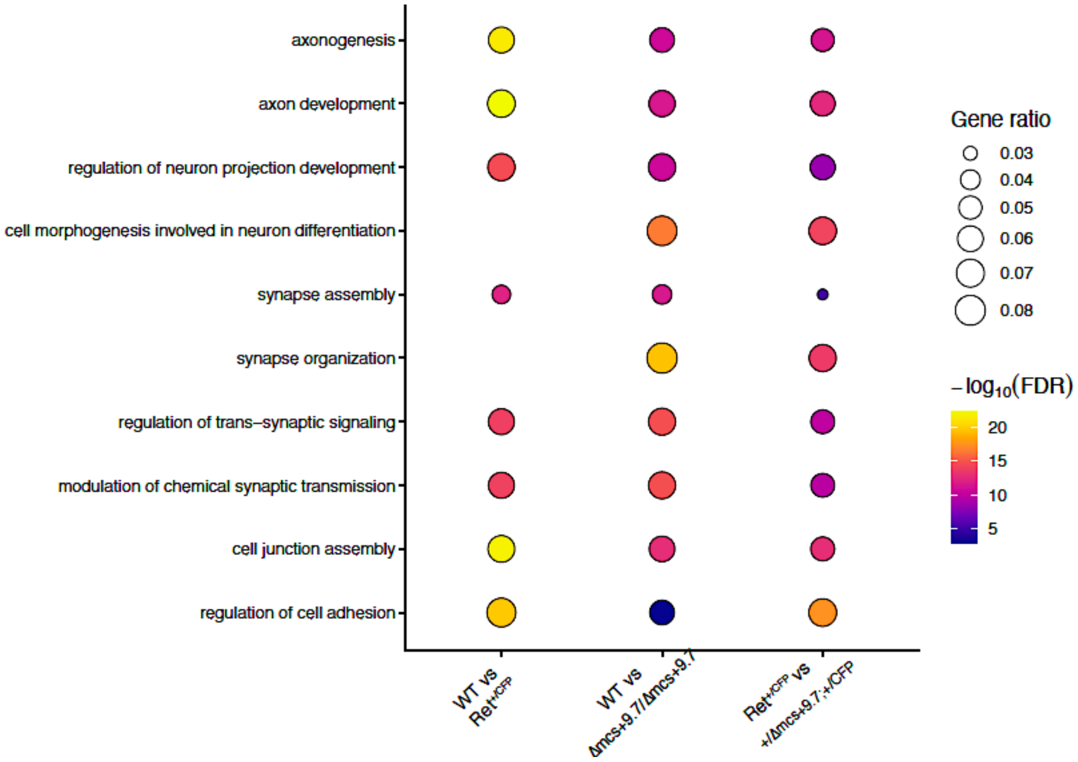


**Supplemental Figure 8:** Bubble plot showing Gene Ontology (GO) enrichment analysis of differentially expressed genes from the three pairwise comparisons: wildtype versus Ret+/CFP, wildtype versus Δmcs+9.7/Δmcs+9.7, and Ret+/CFP versus +/Δmcs+9.7;+/CFP ENS cells. Despite differences in the individual DEGs identified in each comparison, all three datasets converged on a shared set of neuronal developmental processes, including axonogenesis, axon development, regulation of neuron projection development, cell morphogenesis involved in neuronal differentiation, synapse assembly and organization, regulation of trans-synaptic signaling, modulation of chemical synaptic transmission, cell junction assembly, and regulation of cell adhesion. Bubble size indicates the gene ratio, and color denotes enrichment significance (−log10 FDR).

#### Supplemental Figure 9

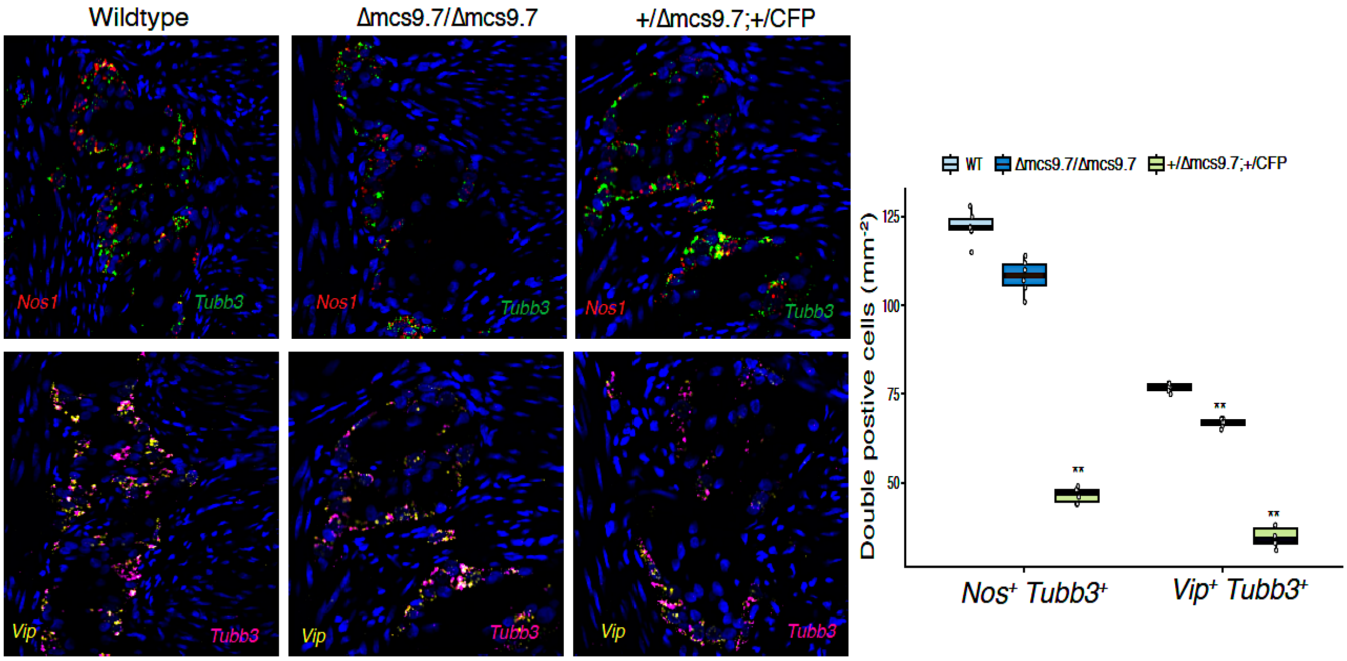


**Supplemental Figure 9:** Representative RNAscope images of E14.5 distal hindgut sections from wildtype, Δmcs+9.7/Δmcs+9.7, and +/Δmcs+9.7;+/CFP embryos stained for *Nos1* (red) and *Tubb3* (green) (top), or *Vip* (yellow) and *Tubb3* (magenta) (bottom). Transcript puncta were assigned to individual segmented cells using CellProfiler, and cells containing both markers were classified as *Nos1*$^+$/*Tubb3*$^+$ or *Vip*$^+$/*Tubb3*$^+$ neurons. Box plots show the density of double-positive neurons (cells $mm^{-2}$) quantified from matched regions of interest. Δmcs+9.7/Δmcs+9.7 embryos exhibited modest reductions in *Nos1*$^+$/*Tubb3* and *Vip*$^+$/*Tubb3*$^+$ neurons relative to wildtype, whereas +/Δmcs+9.7;+/CFP embryos showed markedly greater reductions (61.9% and 54.9%, respectively; $P < 1 \times 10^{-4}$ for both comparisons). Statistical significance was determined using Welch's two-sample *t*-test. Each point represents an independently quantified region of interest
